# Conservation and flexibility in the gene regulatory landscape of Heliconiine butterfly wings

**DOI:** 10.1101/541599

**Authors:** Joseph J Hanly, Richard WR Wallbank, W Owen McMillan, Chris D Jiggins

**Affiliations:** Department of Zoology, University of Cambridge, Downing St., Cambridge CB2 3EJ, UK; Smithsonian Tropical Research Institute, Gamboa, Panama; The George Washington University, Washington DC, 20052, USA

**Keywords:** *cis*-regulation, *Heliconius*, butterfly, transcription factor, homothorax, gene expression, Wnt signaling, transcriptomics

## Abstract

**Background:** Many traits evolve by *cis*-regulatory modification, by which changes to non-coding sequences affect the binding affinity for available transcription factors and thus modify the expression profile of genes. Multiple examples of *cis*-regulatory evolution have been described at pattern switch genes responsible for butterfly wing pattern polymorphism, including in the diverse neotropical genus *Heliconius*, but the identities of the factors that can regulate these switch genes have not been identified.

**Results:** We investigated the spatial transcriptomic landscape across the wings of three closely related butterfly species, two of which have a convergently-evolved, co-mimetic pattern, the other having a divergent pattern. We identified candidate factors for regulating the expression of wing patterning genes, including transcription factors with a conserved expression profile in all three species, and others, including both transcription factors and Wnt pathway genes, with markedly different profiles in each of the three species. We verified the conserved expression profile of the transcription factor homothorax by immunofluorescence, and showed that its expression profile strongly correlates with that of the selector gene optix in butterflies with the Amazonian forewing pattern element ‘dennis’.

**Conclusions:** Here we show that, in addition to factors with conserved expression profiles like homothorax, there are also a variety of transcription factors and signaling pathway components that appear to vary in their expression profiles between closely related butterfly species, highlighting the importance of genome-wide regulatory evolution between species.

## Background

A major challenge in evolutionary developmental biology is to understand how modifications to gene expression can lead to biological diversity. In particular, modulation of *cis*-regulatory elements has been repeatedly identified as the material source of polymorphism and divergence in physiology, behaviour, pigmentation patterns and morphological structures; Gephebase, a database of genotype-phenotype relationships, identifies 323 such examples of *cis*-regulatory evolution in diverse eukaryote clades (Martin and Orgogozo, 2013). These elements must function by differential binding of regulatory factors, and so to understand the evolution of gene regulatory networks, we must first identify which regulatory factors are present and able to perform this function in a given spatial and temporal context.

The Lepidoptera make up ~18% of described animal diversity and have a vast array of wing patterns, both within and between species. Recent advances in genetics, genomics and experimental methods have begun to uncover the underlying genetic and developmental basis of lepidopteran wing pattern variation (Jiggins et al., 2017, Livraghi et al., 2017). One of the most diverse and well-studied groups are the *Heliconius* butterflies, and it is now understood that much of the variation in wing pattern in this group results from regulatory evolution at just three genes, *optix, WntA* and *cortex* (Reed et al., 2011, Martin et al., 2012, Nadeau et al., 2016). CRISPR/Cas9 mutagenesis has shown that *optix* and *WntA* are also involved in the patterning of wings in multiple butterfly lineages (Mazo-Vargas et al., 2017, Zhang et al., 2017). At each of these loci there is a large diversity of complex regulatory alleles, controlling expression patterns across the wing surface during development (Van Belleghem et al., 2017, Enciso-Romero et al., 2017, Wallbank et al., 2016). This regulatory diversity that generates the extraordinary variation in wing patterns nonetheless acts against a background of a highly conserved regulatory factors that underlie insect wing development. Consistent with this idea, earlier candidate gene studies have shown that many patterning factors previously identified in *Drosophila* wing development show similar patterns of expression on butterfly wings (Table 1).

**Table 1.** A summary of previous single gene expression studies in butterfly wing development.

| a) Gene | Species | Reference |
| --- | --- | --- |
| *Ultrabithorax* | *J. coenia* | (Warren et al., 1994, Weatherbee et al., 1999) |
| *apterous* | *J. coenia*, *B. anynana* | (Carroll et al., 1994, Prakash and Monteiro, 2017) |
| *invected* | *J. coenia* | (Carroll et al., 1994) |
| *engrailed* | *P. rapae*, *B. anynana*, *Saturnia pavonia*, *Antheraea polyphemus* | (Monteiro et al., 2006) |
| *scalloped* | *J. coenia* | (Carroll et al., 1994) |
| *wingless* | *V. cardui*, *A. vanillae*, *J. coenia*, *Battus philenor*, *B. mori*, *M. sexta*, *Z. morio*, *B. anynana*, *P. rapae* | (Macdonald et al., 2010, Monteiro et al., 2006) |
| *cut* | *V. cardui*, *A. vanillae*, *J. coenia*, *Battus philenor*, *B. mori*, *M. sexta*, *Z. morio* | (Macdonald et al., 2010) |
| *hedgehog* | *J. coenia* | (Keys et al., 1999) |
| *cubitus interruptus* | *J. coenia* | (Keys et al., 1999) |
| *patched* | *J. coenia* | (Keys et al., 1999) |
| *achaete-scute* | *J. coenia* | (Galant et al., 1998) |
| *Notch* | *H. erato*, *H. melpomene*, *A. vanillae*, *P. rapae*, *M. sexta*, *V. cardui*, *J. coenia*, *B. anynana* | (Reed and Serfas, 2004, Reed, 2004) |
| *Optix* | *Heliconius* spp., *J. coenia*, *V. cardui*, | (Reed et al., 2011, Martin et al., 2014) |
| *WntA* | *Heliconius* spp., *Limenitis arthemis*, *A. vanillae*, *J. coenia*, | (Martin and Reed, 2014, Gallant et al., 2014) |
| *doublesex* | *Papilio* | (Kunte et al., 2014) |
| *aristaless1* & *aristaless2* | *J. coenia*, *Limenitis arthemis*, *Spodoptera ornithogalli*, *Ephestia kuehniella* | (Martin and Reed, 2010) |
| *Distal-less* | *H. erato*, *H. melpomene*, *A. vanillae*, *P. rapae*, *M. sexta*, *V. cardui*, *J. coenia*, *B. anynana*, *Saturnia pavonia*, *Antheraea polyphemus* | (Reed and Serfas, 2004) |
| *spalt major* | *V. cardui*, *P. rapae*, *P. oleracea*, *Colias philodice*, *C. eurytheme*, *B. anynana* | (Zhang and Reed, 2016, Monteiro et al., 2006, Stoehr et al., 2013) |
| *decapentaplegic* | *J. coenia* | (Carroll et al., 1994) |
| *Ecdysone receptor* | *J. coenia* | (Koch et al., 2003) |
| *pSmad* | *B. anynana*, *P. rapae* | (Monteiro et al., 2006) |
| *Antennapedia* | *B. anynana*, *Heteropsis iboina*, *Pararge aegeria* and *Melanargia galathea*, *Lasiommata megera*, *J. coenia*, *M. cinxia*, *Inachis io*, *Caligo memnon* | (Saenko et al., 2011) |
| *cortex* | *H. melpomene*, *H. numata* | (Nadeau et al., 2016) |
| *armadillo* | *B. anynana* | (Connahs et al Biorxiv 2017) |
| *BarH1* | *Colias* | (Woronik et al Biorxiv 2018) |

If this is in fact the case, then the pattern variation we observe in *Heliconius* and other butterflies could be generated by the differential “readout” of a highly conserved set of transcription factors that effectively “prepattern” the wing (figure 1A). These conserved expression patterns could then provide input to the regulatory elements of pattern switch genes like *optix* (figure 1B), and in turn, modifications of these elements could lead to production of wing pattern diversity (figure 1C), a mechanism that allows for the gain of novel phenotypes with the avoidance of deleterious pleiotropic effects (Prud’homme et al., 2007). This hypothesis has previously been examined at the within-species level in *Heliconius* through transcriptomics (Hines et al., 2012) Key examples of this mode of regulatory evolution have been described in evolution of melanic patterns in *Drosophila* species. For example, the protein Engrailed is a deeply conserved component that specifies the posterior compartment in arthropod segmentation (Patel et al., 1989a, Patel et al., 1989b). Some *Drosophila* species have a melanic spot on the anterior tip of the wing, which is sculpted in part by repression of the *yellow* gene by *en*, on the posterior boundary (Gompel et al., 2005). Here, *cis*-regulatory evolution at the *yellow* locus locked onto the conserved spatial information encoded by en. The expression and function of the *en* gene did not change, rather a new regulatory connection was established that modified the expression of yellow to generate novel diversity. Likewise, expression of pigmentation genes in the thorax of *D. melanogaster* is repressed by the expression of the transcription factor *stripe*, which specifies flight muscle attachment sites. The shape of this thoracic element is thus constrained by factors that specify flight muscle pattern (Gibert et al., 2018). On the other hand, there are also examples of *Drosophila* pigmentation evolving by modification to the *trans*-regulatory landscape, by which conserved components of the regulatory landscape themselves change in expression profile to affect downstream melanin synthesis domains. For example; *Dll* in *D. biarmipes* and *D. prolongata* has gained additional expression profiles to its usual peripheral pattern, in correlation with melanic elements (Arnoult et al., 2013). This mode of *trans*-regulatory divergence could also play a role in the evolution of butterfly wing patterns (Figure 1D). No evidence has been found for this at the intraspecific level, but could occur between more distantly related species for which genetic mapping is not possible.

**Figure 1.**
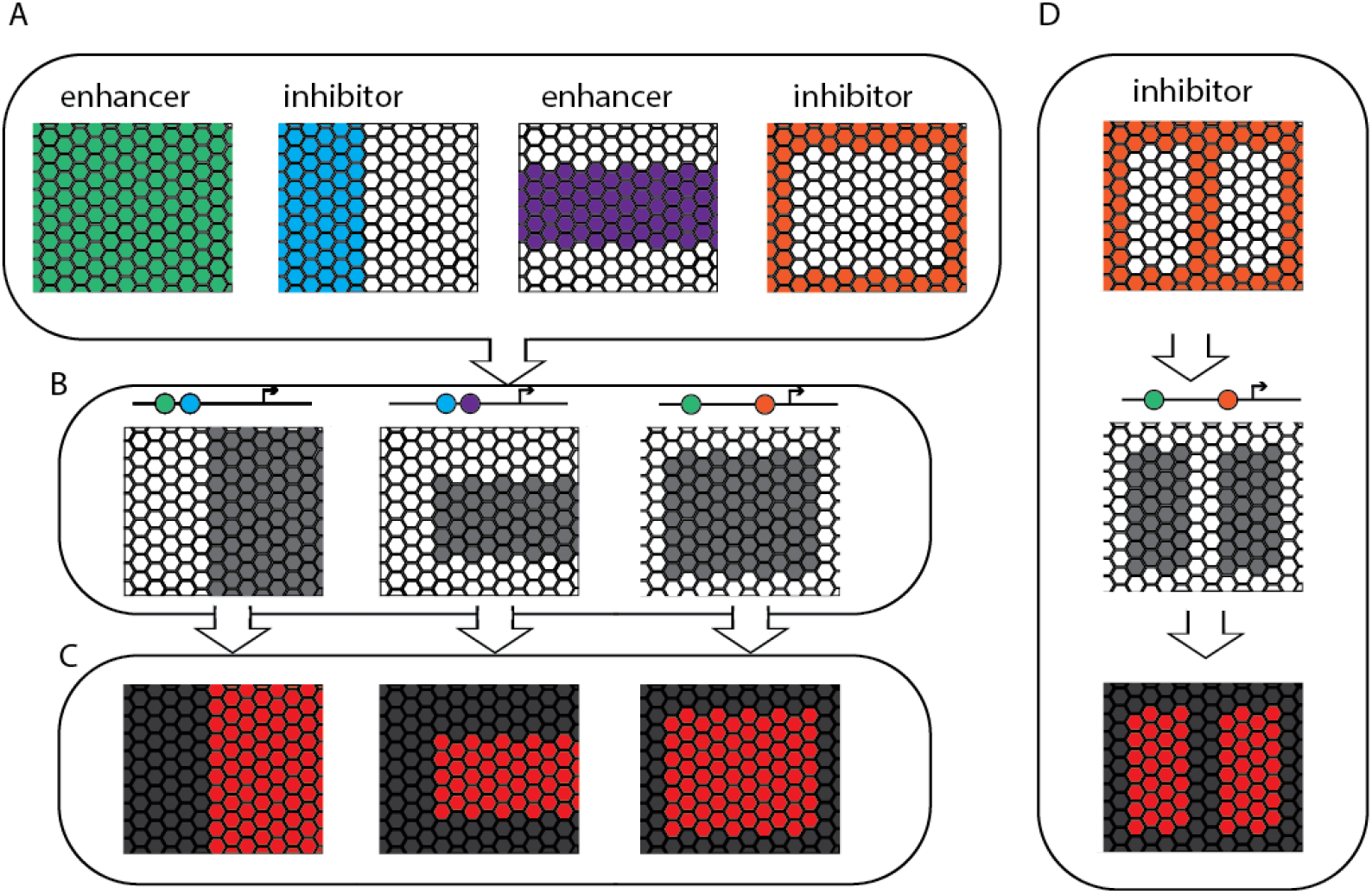
Hypothetical mechanisms of wing pattern development and evolution. In this model, a set of prepatterning factors (A) are expressed early in the developing wing, may pattern the development of the wing, and do not vary in their expression profiles in different morphs or species (see Table 1). These factors feed in to the regulation of the wing pattern switch genes and shape their expression profiles accordingly, for example in *Heliconius* the transcription factor optix (B), which causes scale cells that would otherwise develop to be melanic to express ommochrome pigments (C). It is also possible that changes to the expression of wing pattern switch genes like optix could be caused by changes in expression of prepatterning factors (D).

Here, we begin to test these ideas by using comparative transcriptomic sequencing in two well-described species in the genus *Heliconius*, as well as an outgroup species *Agraulis vanillae*. The two *Heliconius, H. erato* and *H. melpomene*, are co-mimics that diverged from each other between 10-12 mya (Kozak et al., 2015), and recently co-diversified into around 25 different co-occurring wing pattern types. In contrast, *A. vanillae*, which diverged from *Heliconius* roughly 25 mya, is largely monomorphic across its extensive geographic distribution. Linkage and association studies of pattern variation in *Heliconius* and other butterflies have repeatedly identified non-coding regions as primary candidates for the loci of evolution (Kunte et al., 2014, Iijima et al., 2018, Gallant et al., 2014, Wallbank et al., 2016, Van Belleghem et al., 2017). We hypothesize that these candidate regulatory elements allow for regulatory coupling to upstream patterning transcription factors in the wing. In order to understand the upstream spatial information that provides an input to butterfly wing patterning, we need to understand spatial patterns of gene expression in the developing wing, building on a primarily gene-by-gene, candidate driven approach, as in many previous studies which have primarily used factors known from *Drosophila* wing development (Table 1). In particular, these data help us determine which transcription factors show consistent spatial expression profiles in different species and are therefore candidate constituents of a conserved developmental landscape, and which transcription factors show variable patterns and are therefore candidates for the causative regulators of pattern differences. In addition, following on from the discovery that WntA is a key patterning gene in *Heliconius* and *Agraulis* (Mazo-Vargas et al 2017), we also characterised the expression of *Wnt* pathway constituents in all three species. Our results highlight both strong conservation and striking flexibility in the gene regulatory landscape in the early wing development in *Heliconius*.

## Results

To gain insights into the regulatory landscape that organizes butterfly wing early patterning, we conducted transcriptomic analysis of 110 samples representing whole larval wings from *H. melpomene* and *H. erato*, and pupal wings dissected into 5 sections from *H. melpomene, H. erato* and *A. vanillae*. Between 10-24 million reads were sequenced per sample, and the average percentage of reads per sample that did not map was 11.8% (Table S1), compared to a previous RNAseq study in *H. melpomene* in which more than 50% of reads failed to map (Walters et al., 2015). All samples passed quality controls, and could be included in the differentia expression analyses.

PCA analysis showed clustering of samples within each species by stage (Figure 2). Sample clustering by compartment is clear in Day 1 samples, (Figure S1), and in *Agraulis* and *H. melpomene*, distinct clusters for forewing and hindwing are also present, indicating that there is sufficient detectible differential expression between wing sections to determine spatial expression differences across the wing. At Day 2 clustering by compartment is not evident, and there is some clustering by individual, indicating that there is less differential expression across the wing at this later time point.

**Figure 2:**
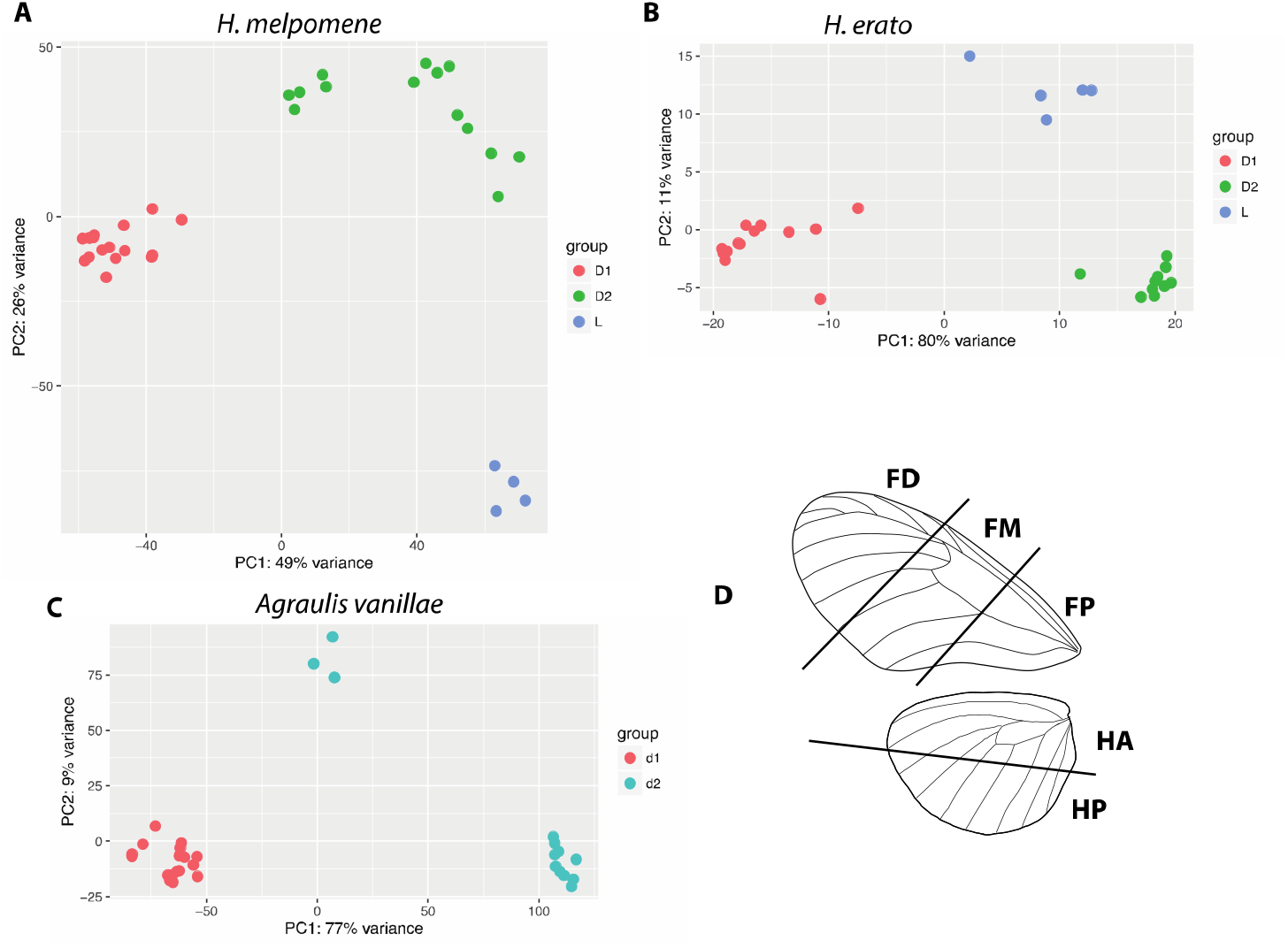
Principal component analyses of RNA samples for each species clustered by stage, with the exception of three samples of *Agraulis vanillae* form the day 2 stage, which formed a separate cluster. D shows the dissection scheme used for tissue collection; FP=Proximal Forewing, FM=Medial Forewing, FD=Distal Forewing, HA=Anterior Hindwing, HP=Posterior Hindwing.

Differential expression analysis revealed 209 differentially expressed genes between *H. melpomene* larval forewings and hindwings, versus 77 in *H. erato*. In total, 28 of these genes are differentially expressed in both species (Table S2). This includes the transcription factor *Ubx*, the notch pathway repressor and microtubule binding protein *pigs*, and 9 genes with no homology to known transcripts. At day 1, we identified 2848 genes differentially expressed between the five compartments in *H. melpomene*, 1713 in *H. erato* and 1780 in *A. vanillae*. 617 transcripts were differentially expressed in all three species at day 1 (Table S3). At day 2, 319 transcripts were differentially expressed in *H. melpomene*, 2663 in *H. erato* and 167 in *A. vanillae*, with no genes differentially expressed in all three species, and 30 genes differentially expressed in a pair of species (Table S4), including the pigmentation genes *Ddc* and *tan*.

### Transcription factors

In order to investigate the nature of the regulatory landscape of the developing wing, we focused on the expression of the 237 identified transcription factor orthology groups. No expression was detected in any species for 37 of these genes; of the 200 TFs that were expressed, 6 were differentially expressed in all three species, 16 were differentially expressed in 2 species, and an additional 31 were differentially expressed in one species (Figure 3). This confirms the presence of TFs that are expressed in a patterned way across the proximal-distal axis of the wing.

**Figure 3:**
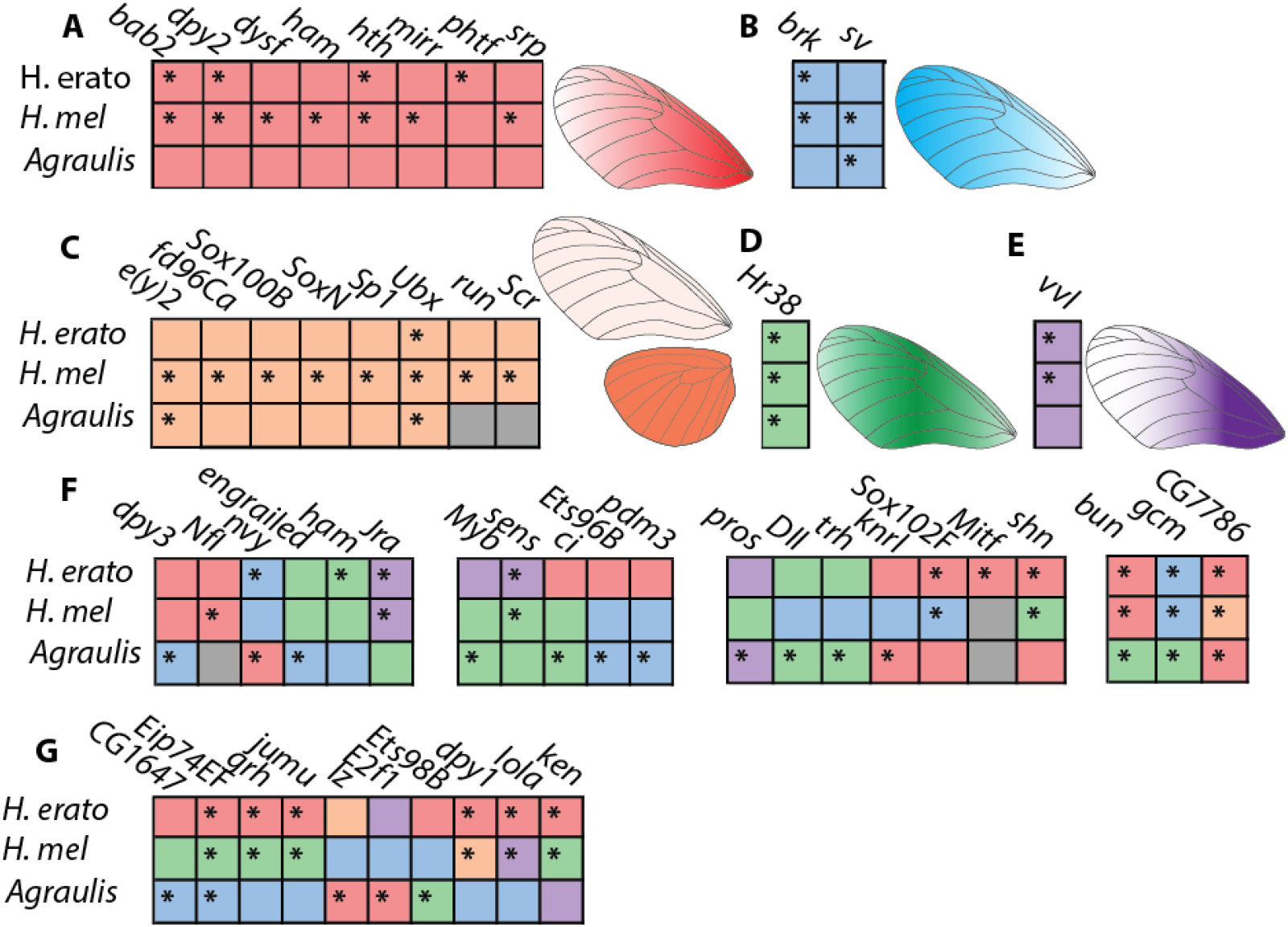
Differential expression of transcription factors in day 1 pupae. Transcription factors are colour coded for their pattern of differential expression – A, in red, indicates factors that are highly expressed in the proximal forewing and expressed in a falling gradient in the medial and distal forewing; B, in blue, indicates factors that are highly expressed in the distal forewing; C, in orange, indicates factors that are highly expressed in the hindwing relative to the forewing; D, in green, indicates factors that are highly expressed in the medial forewing; and E, in purple, indicates factors that are highly expressed in the proximal forewing but low in the rest of the forewing. Asterisks indicate genes which are significantly differentially expressed; all depicted genes are differentially expressed in at least one species.

To examine the relationships between spatial domains of expression between the species, expression profiles were clustered into 5 classes by similarity. A total of 20 TFs shared the same expression profile in all three species and an additional 21 shared the same expression profile in two species, with an additional 10 factors showing a different expression profile in all three species. Of the 37 factors which were differentially expressed in either *H. melpomene* or *H. erato*, 26 (70%) were expressed in the same pattern, whereas of the 27 factors differentially expressed in *Agraulis*, only 13 TFs (48%) were differentially expressed in the same pattern between *Agraulis* and one species of *Heliconius*, indicating that while some factors have conserved ancestral expression profiles, others vary in their expression along the proximal-distal axis.

A number of TFs have expression profiles that match known profiles either from immunohistochemistry of butterfly wings or by analogy with gene expression in *Drosophila* wings (Table 1). This includes *Ultrabithorax* (*Ubx*), expressed only in the hindwing; *homothorax (hth)*, expressed only in the proximal forewing and anterior hindwing; *distal-less (dll)*, expressed in an increasing gradient from proximal to distal; and *mirror (mirr)*, expressed in the proximal forewing and anterior hindwing (Figure 3, c.f. Table 1). The recapitulation of these expression profiles confirms the presence of conserved expression of genes involved with wing pattern specification between insects, and also serves as validation that our experimental design can detect differential expression of transcription factors, which are typically expressed at relatively low levels.

Multiple additional factors with conserved expression profiles along the proximal-distal axis were identified, including *brinker*, a negative-regulator of Dpp signaling (Campbell and Tomlinson, 1999); *bric a brac 2 (bab2)*, part of a proximal-distal gene regulatory module linked to abdominal pigmentation pattern in *Drosophila* (Rogers et al., 2013); *ventral veins lacking (vvl)*, linked to vein development in *Drosophila*, here highly expressed in the proximal forewing, and previously suggested as a candidate wing pattern regulator gene in *H. erato* (Van Belleghem et al., 2017, de Celis et al., 1995); *Hr38*, a hormone receptor upregulated in the medial forewing; and *shaven* (sv), related to the development of sensory structures and upregulated in the distal forewing (Kavaler et al., 1999). These factors serve as additional candidate pre-pattern regulators in heliconiine wings.

Several transcription factors are differentially expressed in all three species, but with different profiles. *bunched (bun)* (Dpp pathway, (Treisman et al., 1995)), *glial cells missing (gcm)* (related to neurogenesis, (Jones et al., 1995)) and *jun-related antigen* (a transcription factor in the JNK-pathway, (Kockel et al., 1997)) are expressed in the same profiles in *Heliconius* but in a different profile in *Agraulis*, whereas the functionally undescribed TF *CG7786* is expressed in a similar profile in *H. erato* and *Agraulis* but differently in *H. melpomene*, and the Wnt-pathway component factors *sens* and *Sox102F* are differentially expressed in different patterns in *H. erato* and *H. melpomene*. In all three species, Ecdysone-induced protein 74EF *Eip74EF* is differentially expressed in a different pattern (Urness and Thummel, 1995). The divergent expression of these factors in this set of species may indicate a role downstream of the pre-patterning factors, for example as targets of Wnt signaling or in scale cell differentiation.

Several differentially expressed TFs are associated with the development of imaginal discs generally, or are specifically associated with other imaginal discs, in particular related to the eye and the genitals suggesting the possible evolution of novel functions in the Lepidoptera. For example *bunched (bun*, eye development), *lozenge (lz*, compound eye development and genital morphogenesis), *ken and barbie* (*ken*, genital morphogenesis). Many differentially expressed TFs have known roles in neurogenesis and the nervous system, including *senseless (sens), gcm, nervy (nvy*, axon guidance and chetae morphogenesis). Other factors have specific associations with cuticle or bristle development such as *nvy, grainy-head (grh)*, as well as multiple copies of *dumpy (dpy)*. The factor *pdm3* is upregulated in *Agraulis* distal hindwing, and is a hotspot for abdominal pigmentation evolution in *Drosophila* (Yassin et al., 2016). This may implicate a novel or undescribed role for these factors in the pupal development of insect wings, or specifically in the wings of butterflies.

### homothorax

The gene *homothorax (hth)* was upregulated in the proximal forewing in all three species. This replicates its previously detected expression in early pupal wings of *H. erato* (Hines et al., 2012). To confirm patterning of the hth protein, pupal butterfly wings were fluorescently stained against an anti-hth antibody raised against *Drosophila* hth. The DNA binding homeodomain of *Heliconius, Danaus* (Monarch butterfly), *Tribolium* and *Drosophila* hth are highly conserved (Figure S3). Staining with anti-Hth highlighted a gradient of hth from the basal to the medial region of the wing in *H. melpomene*. Expression was most strongly detected in presumptive scale cell nuclei (Figure 4)

**Figure 4:**
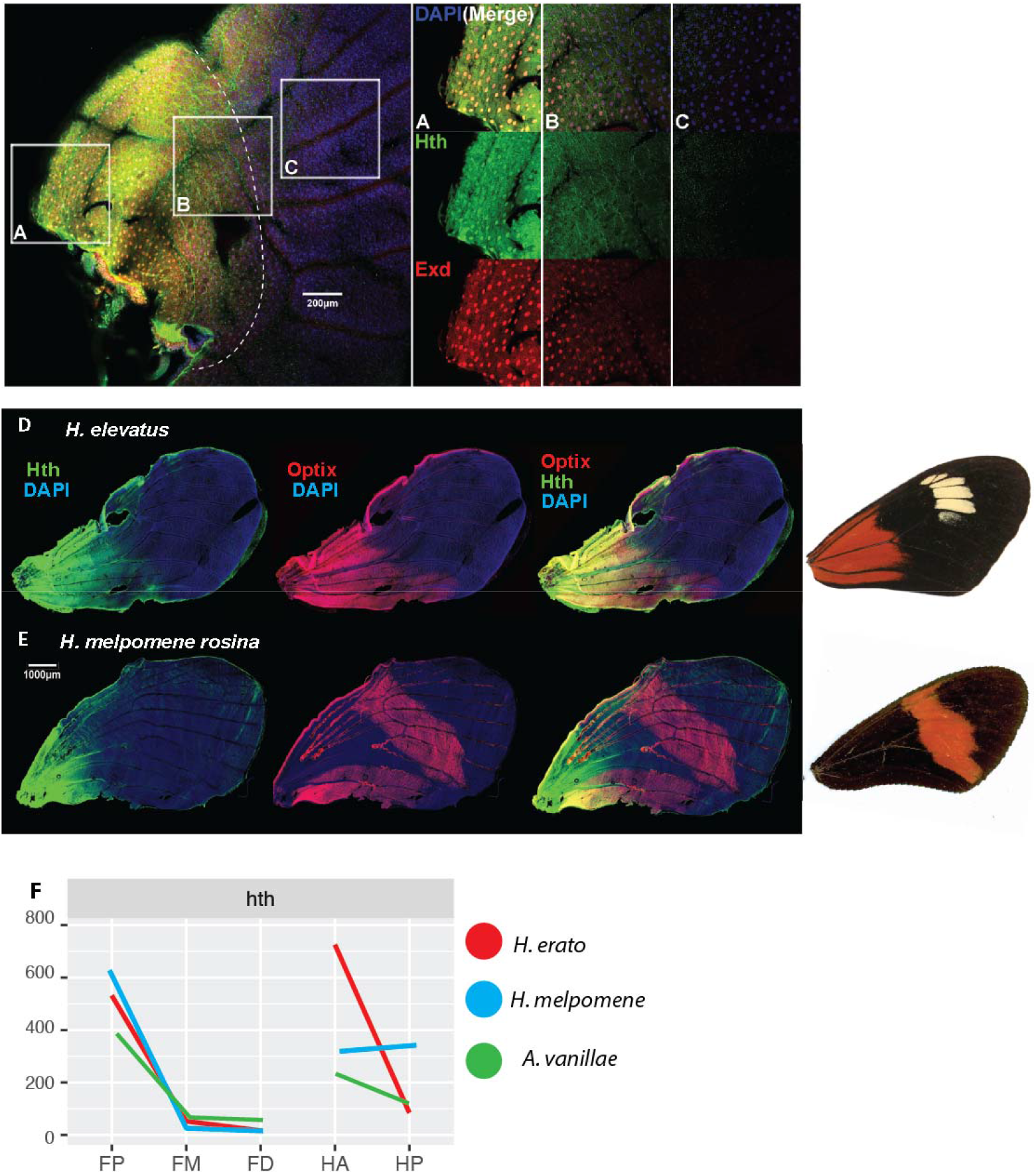
Immunohistochemistry shows pattern of Hth expression in the butterfly wing is replicated by RNAseq analysis. Immunohistochemistry confirmed the expression of Homothorax in a proximal-distal gradient across the basal third of the *Heliconius* wing, in larvae (A-C) and pupae (D, E) of *Heliconius* butterflies. A-C highlight three regions along the proximal-distal axis of the larval wing, showing coincident expression of Homothorax and its co-factor Extradenticle. D shows Homothorax expression in a region coincident with the expression of Optix in a *dennis-ray* butterfly, *Heliconius elevatus*. The same expression pattern of Hth is conserved in a red-banded butterfly (E), but is not associated with Optix expression. All *Heliconius* show Optix expression in the overlapping fore and hindwing region, associated with wing coupling scales as documented previously (Martin et al, 2014). The expression profile observed in pupal wings here recapitulates the levels of *hth* transcript observed in the RNAseq analysis of all three species examined here (F).

Some *Heliconius* wing patterns show a red patch in the proximal forewing that correlates with this *hth* expression domain, known as the *dennis* patch. This patch of red scales is known to be specified by the *optix* gene, and we therefore hypothesised that the evolution of this patch might have arisen through a novel regulatory link between *hth* and *optix*. To explore this possibility, we co-stained wings for both *hth* and *optix* from taxa both with and without the *dennis* patch. In *H. elevatus* which has the *dennis* phenotype, hth expression was strongly coincident with Optix. In contrast in *H. melpomene rosina* which has a medial red forewing band, there was no coincident expression of hth with Optix, except in the region of forewing and hindwing overlap where Optix expression is known to be widely conserved among butterflies. Importantly, *hth* expression was detected from the start of pupation up to and during optix expression at 12-60 hours post-pupation. Together the coincident timing, position and nuclear staining suggest that hth is a potential interacting factor of the *dennis* enhancer in *Heliconius* butterflies. hth was not coincident with the *dennis* bar in the hindwing, however, suggesting other factors are also involved (Figure S4).

### Wnt pathway

Recent studies have highlighted the importance of the ligand WntA in butterfly wing patterning, so we next focused on the expression domains of other Wnt pathway constituents. We identified 52 Wnt pathway constituents in the genomes of our three species (Table S5). Expression profiles were split into three groups based on similarity. In *H. erato* pupal wings, most Wnt pathway genes showed a very similar pattern of highest expression in the proximal forewing and lower expression in the medial and distal forewing, whereas in *H. melpomene* and *A. vanillae*, a variety of different expression profiles for these genes were observed (Figure 5A). The variance in Wnt pathway gene expression is reflected in the prototypical Wnt target transcription factor *senseless* (Figure 3). In contrast to the transcription factors, of the 32 genes differentially expressed at day 1, only 8 (25%) had shared expression profiles in all three species, with an additional 4 (12.5%) sharing a profile between the *Heliconius* species. At day 2, further divergence in expression profiles between species occurred, with none of 19 differentially expressed genes sharing an expression profile between all three species.

**Figure 5:**
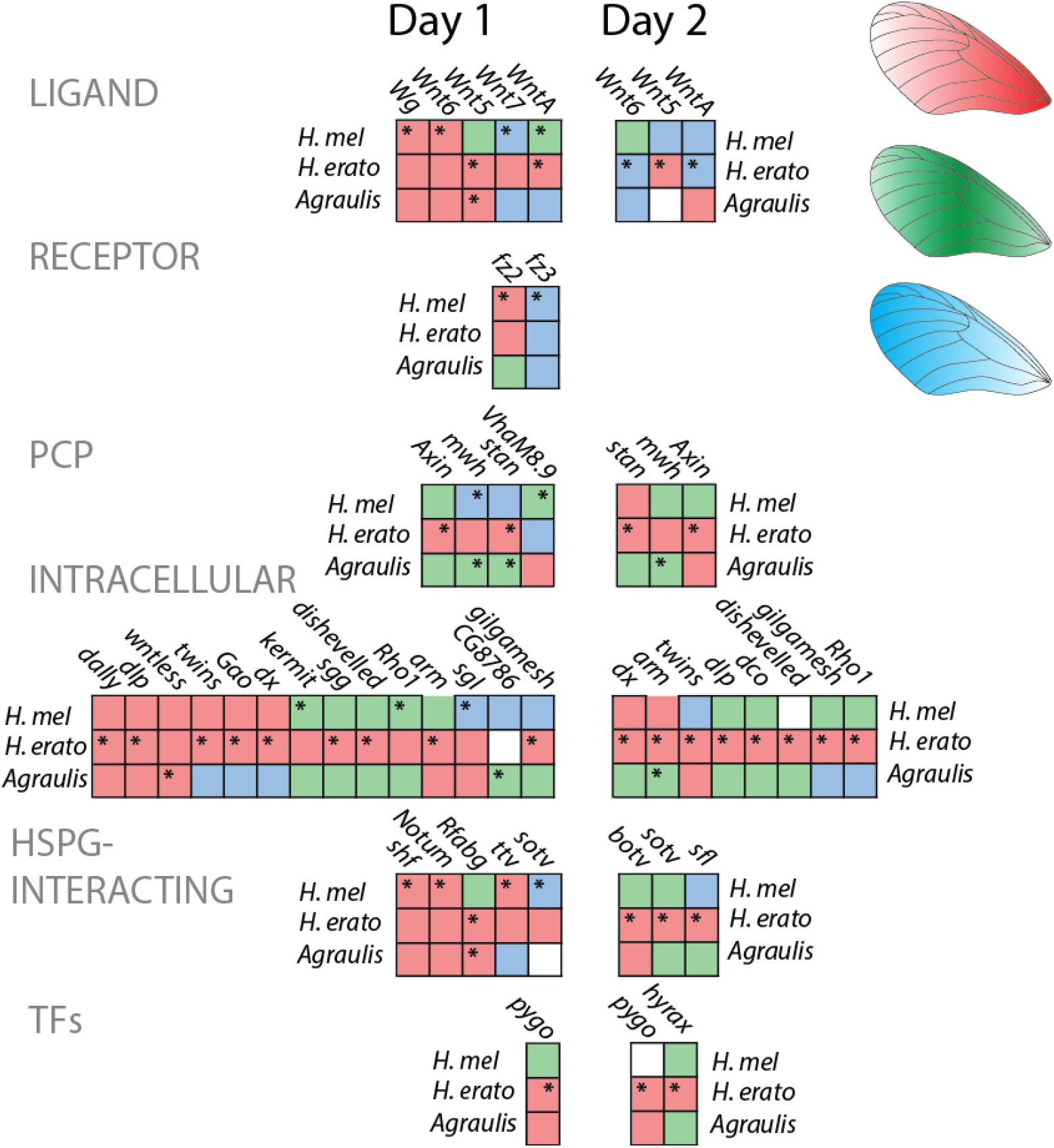
Differential expression of Wnt pathway components in pupal development. Wnt pathway components are colour-coded for their pattern of expression. Red indicates transcripts that were highly expressed in the proximal forewing, green indicates transcripts that were highly expressed in the medial forewing and blue indicates transcripts that were highly expressed in the distal forewing. Asterisks indicate transcripts that were identified as significantly differentially expressed. Note the low discordance of expression profile between species, in contrast to the transcription factors indicated in Figure 3. HSPG = heparin sulfate proteoglycan

The ligand WntA was differentially expressed in the *Heliconius* species in profiles that correlate to that observed in Martin et al (2012), and multiple other Wnt ligands were differentially expressed in the three species, including Wnt2 and Wnt6 in *Heliconius*, which in other Nymphalid species have redundant expression profiles and have been correlated with the discalis pattern elements, and which was also previously reported in *H. erato* (Martin et al., 2014, Hines et al., 2012). Two Wnt receptors, *fz2* and *fz3*, are expressed in opposition to each other; fz2 is the primary Wg receptor in *Drosophila*, while fz3 plays an inhibitory role (Bhanot et al., 1996, Sato et al., 1999). Wnt pathway components involved with planar cell polarity including *multiple wing hairs (mwh), starry night (stan)*, and *Axin*, were differentially expressed at both stages, along with other intracellular components of the Wnt pathway including *armadillo*/β-*catenin* and *wntless*, a transmembrane factor required for Wnt ligand secretion. In *H. erato*, two TFS downstream of Wnt signaling were differentially expressed; *pygo* and *hyrax*.

## Discussion

We have explored the patterns of gene expression both through development, across evolutionary divergence and across the developing wing. Our results paint a vivid picture of how wing patterns develop and evolve across the three butterfly species. Broadly, *H. melpomene, H. erato* and *A. vanillae* share a common spatial transcriptomic landscape in the developing wing with other Lepidoptera and with *Drosophila*, implying the existence of a shared insect wing gene regulatory network (cf. Table 1). The results for *H. erato* here also broadly replicate those from a previous transcriptomic analysis there(Hines et al., 2012). However, our data also highlight considerable flexibility in the transcriptional landscape. More than half of the transcription factors that are differentially expressed across the developing wing surface have different expression profiles within the heliconiines. This flexibility is most evident within the Wnt signaling pathway constituents, where *H. erato* has a derived pattern of strongly coordinated Wnt pathway gene expression, that is not seen in the other two species, despite strong convergence in pattern between the two *Heliconius* species studied. Recent work has shown genome-wide selection on regulatory elements at the between-population level in *Heliconius (Lewis and Reed, 2018)*, and it is likely that in the ancestral linages of each species, many functional changes could be accrued that would lead to many differences in patterns of gene expression in the wing.

### Transcription factors expressed in butterfly wings

Transcription factors provide the physical interactions that lead to differential regulation in gene regulatory networks. Several transcription factors that are known to be involved in development of wings in *Drosophila* and *Junonia* were identified in this experiment in their expected expression profiles, including *Ubx* and *hth*. Several other factors were expressed in similar patterns in all three species in this experiment; these additional factors could delineate the developmental morphospace along the forewing proximal-distal axis and hindwing anteroposterior axis in pupal wings. We identified an additional cohort of transcription factors with non-conserved expression profiles between the three studied species; many more of these factors had shared expression profiles between *H. melpomene* and *erato* than between *Agraulis* and Heliconius. The two Heliconius species are more closely related to one another, but are also convergent in their wing patterns, so we cannot currently disentangle whether the share expression patterns are due to common ancestry or are convergent due to shared selection pressures. Others were different in all three species, implying developmental drift, or a lack of constraint, on the regulation of these factors. Such factors have the potential to act as the substrate for functional diversification.

### hth implicated in mimetic pattern evolution

One of the most strongly and consistently differentiated transcription factors was hth, and we therefore followed up on the expression patterns using immunohistochemistry. This confirmed that the hth protein shows a conserved pattern of expression restricted to the proximal wing region in all species examined. Furthermore, co-staining with anti-optix demonstrated a strong correlation of hth with the expression of optix protein in butterflies with the red proximal *dennis* patch. Expression of both proteins was localized to scale cell nuclei, and spatial patterns were tightly correlated between the two factors. In contrast, butterflies lacking the dennis patch showed a conserved expression of hth, but no correlated expression of optix. hth a candidate regulator of optix in dennis+ butterflies. A possible mechanism for the evolution of the dennis pattern is therefore that an *optix* regulatory region gained transcription factor binding sites for hth, allowing the development of a novel pattern without the requirement for changes to the expression or function of hth itself, similar to the roles of *en* and *sr* in the development of melanic patterns in *Drosophila* (Gompel et al., 2005, Gibert et al., 2018). Alternatively, hth might regulate optix through intermediate factors, and a number of other candidates upregulated in the proximal region are evident from our results. Future analyses of the dennis regulatory element will be required to determine the precise mechanisms of interaction with upstream regulators.

### Wnt pathway variance implies different functions

Variance in *WntA* expression in correlation with wing pattern has previously been shown in many butterfly clades, including between races and species of *Heliconius*, and in *Agraulis* (Martin and Reed, 2014, Martin et al., 2012). We found that other Wnt pathway constituents also vary in their expression domains between species.

Surprisingly, expression patterns of Wnt pathway constituents were completely different between the two *Heliconius* co-mimics. In particular, the Wnt pathway constituents in *H. erato* were mainly expressed in a correlated pattern – high in the proximal forewing, and low everywhere else, whereas the same factors were expressed in a variety of patterns in *Agraulis* and *H. melpomene*.

Notably, the correlated pattern in *H. erato* closely mirrors the expression profile of *WntA* in larval wing discs of *H. erato demophoon* from Panama, the pattern form used in this study (Martin et al., 2012), while the *WntA* expression profile for Panamanian *H. melpomene rosina* is notably different from its co-mimic. Here, expression is present in the distal forewing as well as the proximal forewing, and the boundary of proximal *WntA* expression does not correlate well with the proximal boundary of the red pattern element in the adult wing.

Together, these differences in both WntA function and Wnt pathway component expression could suggest that regulatory diversification of the Wnt pathway has occurred between *Heliconius* species, implying that this aspect of the wing gene regulatory network has diverged in the lineages leading to these mimetic forms, requiring the utilization of different functional mechanisms for building an identical wing pattern. This suggests extensive genome-wide regulatory divergence at the between-species level between *Heliconius* butterflies. Alternatively, the differences in expression of Wnt pathway components could entirely be a consequence of the different expression profiles of WntA between the two species. Wnt ligands in *Drosophila* effect their own autoregulation through modulation of receptor, ligand and transcription factor expression levels (Cadigan et al., 1998, Schilling et al., 2014). If WntA is capable of directly regulating the expression of Wnt pathway components, this could explain all the differences we observed here while requiring between-species regulatory divergence at just one genomic locus. Separating these two models will require the functional exploration of the effects of WntA signaling.

## Conclusion

Our understanding of the regulatory evolution of wing pattern in butterflies is dependent on a clear picture of the expression of developmental factors around the time of wing pattern specification. This study has provided a picture of gene expression along one axis of developing wings in a manner unbiased by our understanding of wing development in non-lepidopteran systems. At the within-species level, we can broadly rule out the hypothesis that *trans*-regulatory factors change their expression profiles in different pattern forms (i.e. Figure 1D) based on genetic mapping, but we are not able to rule out this phenomenon at the between-species level – it is likely that both processes play a role, either through selection or drift. Our deeper understanding of factors that are expressed in the wing in correlation with pattern elements will permit us to decode the regulatory linkages that lead to the differential expression of pattern switch genes like *optix, WntA* and *cortex*, and it is clear that we should look to both conserved and diverging regulatory factors as the causative agents of *cis*-regulatory evolution.

## Methods

### Tissue sampling and dissection

*Heliconius melpomene rosina* and *Heliconius erato demophoon* were collected from stocks maintained at the Smithsonian Tropical Research Institute in Gamboa, Panama in February and July 2014. Adults were provided with an artificial diet of pollen/glucose solution supplemented with flowers of *Psiguria, Lantana* and/or *Psychotria alata* according to availability. Females were provided with *Passiflora* plants for egg laying (P. *menispermifolia* for *H. melpomene, P. biflora* for *H. erato)*. Eggs were collected daily, and caterpillars reared on fresh shoots of *P. williamsi (melpomene)* or *P. biflora (erato)* until late 5th (final) instar, when they were separated into individual pots in a temperature-monitored room, and closely observed for the purpose of accurate developmental staging. *Agraulis vanillae* larvae were collected from *P. edulis* located near the insectary in March 2014.

Pre-pupation larvae were identified for dissection. Late 5th instar larvae of *Heliconius* undergo colour changes from white to purple on the last larval day, followed by an additional change to pink-orange in the hours before pupation. Additionally, several behavioural changes accompany the pre-pupation period; the larvae stop eating and clear their digestive tract, then undertake a period of rapid locomotion and wandering until they find an appropriate perch for pupation - preferably the underside of a leaf or a sturdy twig - at which point they settle in place and produce a strong silk attachment. Gradually, over a period of 30-120 minutes, they suspend themselves form their perch in a J shape and then pupate. Larvae that were post-locomotion but pre-J-shape were dissected in cold PBS and the wing discs removed.

Pupae were allowed to develop until 36h (+/− 1.5h), or to 60h (+/− 1.5h). These time points are referred to as Day 1 and Day 2 throughout. In the hours immediately post-pupation (Day 0), the pupal carapace is soft and the membranous structures of the pupa are thin, weak, transparent and sticky, hence the effective dissection of unfixed, intact pupal structures is very challenging at the earliest pupal time points.

Pupae were dissected in cold PBS. Wings were removed from the pupa and cleared of peripodial membrane. The wings were then cut with microdissection scissors into 5 sections: forewing proximal, medial and distal, and hindwing anterior and posterior (figure 2). The lacunae (developing veins) were used as landmarks for dissection.

Whole larval wing discs and pupal wing sections were transferred into RNAlater (ThermoFisher, Waltham, MA) and kept on liquid nitrogen in Gamboa, then transported to the UK on dry ice, and transferred to −80 C on arrival in Cambridge.

### RNA extraction and sequencing

RNA extraction was carried out using a standard hybrid protocol. Briefly, wing tissue sections were transferred into Trizol Reagent (Invitrogen, Carlsbad, CA) and disassociated using stainless steel beads in a tissue lyser. Chloroform phase extraction was performed, followed by purification with the RNeasy kit (Qiagen, Valencia CA). RNA was eluted into distilled water and treated with DNAseI (Ambion, Naugatuck CT), then quantified and stored at −80 C. Left and right wings and wing sections were pooled.

cDNA synthesis, library preparation and sequencing were carried out by Beijing Genomics Institute (Beijing, China). Samples were sequenced at either 75 PE on Illumina HiSeq 3000 or at 150 PE on Illumina HiSeq 4000.

### Transcriptome assembly - Agraulis

All paired end sequence data for *Agraulis* was assembled with the transcriptome assembler Trinity (Haas et al., 2013). This generated 87,214 contigs. Next the Trinity output was passed through TransDecoder (Haas & Papanicolaou, in prep), which annotates the transcript contigs based on the likelihood that they contain reading frames, and also based on similarity by BLAST of transcripts to reference assemblies, in this case *H. erato* and *H. melpomene*. This annotation (a GFF3 annotation of the Trinity contigs) contained 24,984 genes, which compares to 20,102 annotated genes in *H. melpomene v2.1* and 13,676 in *H. erato v1*.

### Mapping and quantification

Reads were aligned with Hisat2 aligner to the genome of the respective species (Rutledge et al., 2006, Kim et al., 2015). The highest percentage of unique mappings was achieved using default parameters (Figure 3.4). Alignments were then quantified using GFF annotations of each genome with HTSeqCount, union mode (Anders et al., 2015). Genomes and annotations are publicly available at www.lepbase.org (Challis et al., 2016).

### Data analysis

Statistical analysis of counts was carried out using the R package DESeq2 (Love et al., 2014) using the following generalized linear model (GLM):

~ *individual + compartment*

In larvae, the compartments were Forewing (FW) and Hindwing (HW). In pupae, the compartments were as follows: Proximal Forewing (FP), Medial Forewing (FM), Distal Forewing (FD), Anterior Hindwing (HA), Posterior Hindwing (HPo) (figure 2)).

### Determining homology

Homology between differentially expressed genes in the three species was determined in two ways. First, a small percentage of genes have been assigned homologs by comparison with other lepidopteran genomes on LepBase using InterProScan (Jones et al., 2014). This allowed recovery of one-to-one homologs and gene families with distinct insect lineages. However in a number of cases where similar copies of a gene are present, for example the Wnt ligands, some genes were assigned to the incorrect orthogroup and were manually curated. For the rest of the genes, as well as all genes in *Agraulis*, amino acid sequences were reciprocally searched with BLASTp, and the top hit was taken as the homolog (Altschul et al., 1997). Genes with no assigned orthogroup were compared by pBLAST against the polypeptide library of *D. melanogaster* genes retrieved from FlyBase, associating them with a FBpp number and gene code based on homology with *Drosophila* genes (Gramates et al., 2017).

### Immunohistochemistry

Pupae were dissected 60-80 hrs after pupation in chilled PBS, or at an estimated 12hrs before pupation for final instar larvae. Wings were fixed in 4% MeOH-free formaldehyde (Pierce) in PBSTw (PBS plus 0.01% Tween-20 (Sigma)) on ice for 40mins. They were then washed and permeabilized 6 x 5mins in PBSTx (PBS plus 0.5% Triton-X (Sigma)) and blocked for 2 hrs room temperature in PBSTx plus 5% goat serum (Sigma). Rabbit anti-Homothorax antibody (gift from Prof. Adi Salzberg, Technion-Israel Institute of Technology) or Rat anti-Optix (gift from Prof. Robert Reed, Cornell University) antibody was diluted to 1/1000 in PBSTx plus 0.5% goat serum and applied to wings overnight at 4° C. Wings were washed 6×5mins in PBSTx and incubated with 1/1000 goat anti-rabbit alexa-488 conjugated antibody (Abcam) in PBSTx for 3hrs at room temperature. Wings were washed 4×5mins with PBSTx and incubated 10mins in 1ug/ml DAPI (ThermoScientific) PBSTx. Wings were mounted in Fluoromount-G (SouthernBiotech) and imaged using a Leica DM6000B SP5 confocal microscope. An average of 60-80 images were taken to cover the entire wing, each a stack of 40 2 μm thick slices and composed of three channels: 408 nm, 488 nm, and 568 nm. Each stack was converted to single images of maximum intensity using FIJI ImageJ 1.47m, and then split by channel. Images were then compiled into a single image and formatted using Adobe Photoshop CS5.1.

## Declarations

### Data availability

The datasets generated and/or analysed during the current study are available in the SRA repository [PERSISTENT WEB LINK TO DATASETS]

### Ethics Approval and Consent to Participate

Not applicable

### Consent for Publication

Not applicable.

### Author contributions

JJH, WOM and CDJ planned the research and wrote the paper, JJH performed experiments and analysis, RWRW performed immunofluorescence and imaging.

## Acknowledgements

We thank Lucas Brenes, Henry Arenas-Castro, Oscar Paneso and Elizabeth Evans for assistance with insect rearing and tissue dissection, and the Beijing Genomics Institute for library preparation and sequencing. We thank Arnaud Martin for comments on an earlier draft of this text. We thank Adi Salzberg for use of the Hth antibody. We thank the Smithsonian Tropical Research Institute for support for tissue collection and work with butterflies in Panama, and ANAM in Panama for permission to collect butterflies.

## References

Altschul, S. F., Madden, T. L., Schaffer, A. A., Zhang, J., Zhang, Z., Miller, W. & Lipman, D. J. 1997. Gapped BLAST and PSI-BLAST: a new generation of protein database search programs. Nucleic Acids Res, 25, 3389–402.

Anders, S., Pyl, P. T. & Huber, W. 2015. Htseq~A Python Framework To Work With High-Throughput Sequencing Data. Bioinformatics, 31, 166–9.

Arnoult, L., Su, K. F., Manoel, D., Minervino, C., Magrina, J., Gompel, N. & Prud’Homme, B. 2013. Emergence and diversification of fly pigmentation through evolution of a gene regulatory module. Science, 339, 1423–6.

Bhanot, P., Brink, M., Samos, C. H., Hsieh, J. C., Wang, Y., Macke, J. P., Andrew, D., Nathans, J. & Nusse, R. 1996. A new member of the frizzled family from Drosophila functions as a Wingless receptor. Nature, 382, 225–30.

Cadigan, K. M., Fish, M. P., Rulifson, E. J. & Nusse, R. 1998. Wingless repression of Drosophila frizzled 2 expression shapes the Wingless morphogen gradient in the wing. Cell, 93, 767–77.

Campbell, G. & Tomlinson, A. 1999. Transducing the Dpp Morphogen Gradient In the wing of Drosophila: regulation of Dpp targets by brinker. Cell, 96, 553–62.

Carroll, S. B., Gates, J., Keys, D. N., Paddock, S. W., Panganiban, G. E., Selegue, J. E. & Williams, J. A. 1994. Pattern formation and eyespot determination in butterfly wings. Science, 265, 109–14.

Challis, R. J., Kumar, S., Dasmahapatra, K. K. K., Jiggins, C. D. & Blaxter, M. 2016. Lepbase: the Lepidopteran genome database. BioRxiv, 056994.

De Celis, J. F., Llimargas, M. & Casanova, J. 1995. Ventral veinless, the gene encoding the Cf1a transcription factor, links positional information and cell differentiation during embryonic and imaginal development in Drosophila melanogaster. Development, 121, 3405–16.

Enciso-Romero, J., Pardo-Diaz, C., Martin, S. H., Arias, C. F., Linares, M., Mcmillan, W. O., Jiggins, C. D. & Salazar, C. 2017. Evolution of novel mimicry rings facilitated by adaptive introgression in tropical butterflies. Molecular Ecology, 26, 5160–5172.

Galant, R., Skeath, J. B., Paddock, S., Lewis, D. L. & Carroll, S. B. 1998. Expression pattern of a butterfly achaete-scute homolog reveals the homology of butterfly wing scales and insect sensory bristles. CurrBiol, 8, 807–13.

Gallant, J. R., Imhoff, V. E., Martin, A., Savage, W. K., Chamberlain, N. L., Pote, B. L., Peterson, C., Smith, G. E., Evans, B., Reed, R. D., Kronforst, M. R. & Mullen, S. P. 2014. Ancient homology underlies adaptive mimetic diversity across butterflies. Nat Commun, 5, 4817.

Gibert, J. M., Mouchel-Vielh, E. & Peronnet, F. 2018. Pigmentation Pattern And Developmental Constraints: Flight Muscle Attachment sites delimit the thoracic trident of Drosophila melanogaster. Sci Rep, 8, 5328.

Gompel, N., Prud’Homme, B., Wittkopp, P. J., Kassner, V. A. & Carroll, S. B. 2005. Chance caught on the wing: *cis*-regulatory evolution and the origin of pigment patterns in Drosophila. Nature, 433, 481–7.

Gramates, L. S., Marygold, S. J., Santos, G. D., Urbano, J. M., Antonazzo, G., Matthews, B. B., Rey, A. J., Tabone, C. J., Crosby, M. A., Emmert, D. B., Falls, K., Goodman, J. L., Hu, Y., Ponting, L., Schroeder, A. J., Strelets, V. B., Thurmond, J., Zhou, P. & The Flybase, C. 2017. Flybase At 25: looking to the future. Nucleic Acids Res, 45, D663–D671.

Haas, B. J., Papanicolaou, A., Yassour, M., Grabherr, M., Blood, P. D., Bowden, J., Couger, M. B., Eccles, D., Li, B., Lieber, M., Macmanes, M. D., Ott, M., Orvis, J., Pochet, N., Strozzi, F., Weeks, N., Westerman, R., William, T., Dewey, C. N., Henschel, R., Leduc, R. D., Friedman, N. & Regev, A. 2013. De novo transcript sequence reconstruction from RNA-seq using the Trinity platform for reference generation and analysis. Nat Protoc, 8, 1494–512.

Hines, H. M., Papa, R., Ruiz, M., Papanicolaou, A., Wang, C., Nijhout, H. F., Mcmillan, W. O. & Reed, R. D. 2012. Transcriptome analysis reveals novel patterning and pigmentation genes underlying Heliconius butterfly wing pattern variation. BMC Genomics, 13, 288.

Iijima, T., Kajitani, R., Komata, S., Lin, C. P., Sota, T., Itoh, T. & Fujiwara, H. 2018. Parallel Evolution Of Batesian Mimicry Supergene In Two Papilio butterflies, P. polytes and P. memnon. Sci Adv, 4, eaao5416.

Jiggins, C. D., Wallbank, R. W. & Hanly, J. J. 2017. Waiting In The Wings: What Can We Learn about gene co-option from the diversification of butterfly wing patterns? Philos Trans R Soc Lond B Biol Sci, 372.

Jones, B. W., Fetter, R. D., Tear, G. & Goodman, C. S. 1995. glial cells missing: a genetic switch that controls glial versus neuronal fate. Cell, 82, 1013–23.

Jones, P., Binns, D., Chang, H. Y., Fraser, M., Li, W., Mcanulla, C., Mcwilliam, H., Maslen, J., Mitchell, A., Nuka, G., Pesseat, S., Quinn, A. F., Sangrador-Vegas, A., Scheremetjew, M., Yong, S. Y., Lopez, R. & Hunter, S. 2014. InterProScan 5: genome-scale protein function classification. Bioinformatics, 30, 1236–40.

Kavaler, J., Fu, W., Duan, H., Noll, M. & Posakony, J. W. 1999. An essential role for the Drosophila Pax2 homolog in the differentiation of adult sensory organs. Development, 126, 2261–72.

Keys, D. N., Lewis, D. L., Selegue, J. E., Pearson, B. J., Goodrich, L. V., Johnson, R. L., Gates, J., Scott, M. P. & Carroll, S. B. 1999. Recruitment of a hedgehog regulatory circuit in butterfly eyespot evolution. Science, 283, 532–4.

Kim, D., Langmead, B. & Salzberg, S. L. 2015. Hisat: A Fast Spliced Aligner With Low memory requirements. Nat Methods, 12, 357–60.

Koch, P. B., Merk, R., Reinhardt, R. & Weber, P. 2003. Localization of ecdysone receptor protein during colour pattern formation in wings of the butterfly Precis coenia (Lepidoptera: Nymphalidae) and co-expression with Distal-less protein. Dev Genes Evol, 212, 571–84.

Kockel, L., Zeitlinger, J., Staszewski, L. M., Mlodzik, M. & Bohmann, D. 1997. Jun In Drosophila development: redundant and nonredundant functions and regulation by two MAPK signal transduction pathways. Genes Dev, 11, 1748–58.

Kozak, K. M., Wahlberg, N., Neild, A. F., Dasmahapatra, K. K., Mallet, J. & Jiggins, C. D. 2015. Multilocus species trees show the recent adaptive radiation of the mimetic heliconius butterflies. Syst Biol, 64, 505–24.

Kunte, K., Zhang, W., Tenger-Trolander, A., Palmer, D. H., Martin, A., Reed, R. D., Mullen, S. P. & Kronforst, M. R. 2014. doublesex is a mimicry supergene. Nature, 507, 229–32.

Lewis, J. J. & Reed, R. D. 2018. Genome-wide regulatory adaptation shapes population-level genomic landscapes in Heliconius. Mol Biol Evol.

Livraghi, L., Arnaud Martin, Melanie Gibbs, Nora Braak, Saad Arif & Breuker., C. J. 2017. “CRISPR/Cas9 as the Key to Unlocking the Secrets of Butterfly Wing Pattern Development and Its Evolution. “.Advances in Insect Physiology.

Love, M. I., Huber, W. & Anders, S. 2014. Moderated estimation of fold change and dispersion for RNA-seq data with DESeq2. Genome biology, 15, 550.

Macdonald, W. P., Martin, A. & Reed, R. D. 2010. Butterfly wings shaped by a molecular cookie cutter: evolutionary radiation of lepidopteran wing shapes associated with a derived Cut/wingless wing margin boundary system. Evol Dev, 12, 296–304.

Martin, A., Mcculloch, K. J., Patel, N. H., Briscoe, A. D., Gilbert, L. E. & Reed, R. D. 2014. Multiple recent co-options of Optix associated with novel traits in adaptive butterfly wing radiations. Evodevo, 5, 7.

Martin, A. & Orgogozo, V. 2013. The Loci Of Repeated Evolution: A Catalog of genetic hotspots of phenotypic variation. Evolution, 67, 1235–50.

Martin, A., Papa, R., Nadeau, N. J., Hill, R. I., Counterman, B. A., Halder, G., Jiggins, C. D., Kronforst, M. R., Long, A. D., Mcmillan, W. O. & Reed, R. D. 2012. Diversification of complex butterfly wing patterns by repeated regulatory evolution of a Wnt ligand. Proc Natl Acad Sci USA, 109, 126327.

Martin, A. & Reed, R. D. 2010. Wingless And Aristaless2 Define A Developmental Ground Plan For Moth and butterfly wing pattern evolution. Mol Biol Evol, 27, 2864–78.

Martin, A. & Reed, R. D. 2014. Wnt Signaling Underlies Evolution And Development of the butterfly wing pattern symmetry systems. Dev Biol, 395, 367–78.

Mazo-Vargas, A., Concha, C., Livraghi, L., Massardo, D., Wallbank, R. W. R., Zhang, L., Pap Ador, J. D., Martinez-Najera, D., Jiggins, C. D., Kronforst, M. R., Breuker, C. J., Reed, R. D., Patel, N. H., Mcmillan, W. O. & Martin, A. 2017. Macroevolutionary shifts of WntA function potentiate butterfly wing-pattern diversity. Proc Natl Acad Sci USA, 114, 10701–10706.

Monteiro, A., Glaser, G., Stockslager, S., Glansdorp, N. & Ramos, D. 2006. Comparative Insights Into Questions of lepidopteran wing pattern homology. BMC Dev Biol, 6, 52.

Nadeau, N. J., Pardo-Diaz, C., Whibley, A., Supple, M. A., Saenko, S. V., Wallbank, R. W., Wu, G. C., Maro Ja, L., Ferguson, L., Hanly, J. J., Hines, H., Salazar, C., Merrill, R. M., Dowling, A. J., Ffrench-Constant, R. H., Llaurens, V., Joron, M., Mcmillan, W. O. & Jiggins, C. D. 2016. The Gene Cortex Controls Mimicry And Crypsis In Butterflies And Moths. Nature, 534, 106–10.

Patel, N. H., Kornberg, T. B. & Goodman, C. S. 1989a. Expression of engrailed during segmentation in grasshopper and crayfish. Development, 107, 201–12.

Patel, N. H., Martin-Blanco, E., Coleman, K. G., Poole, S. J., Ellis, M. C., Kornberg, T. B. & Goodman, C. S. 1989b. Expression of engrailed proteins In arthropods, annelids, and chordates. Cell, 58, 955–68.

Prakash, A. & Monteiro, A. 2017. apterous A Specifies Dorsal Wing Patterns And Sexual Traits In Butterflies. bioRxiv, 131011.

Prud’Homme, B., Gompel, N. & Carroll, S. B. 2007. Emerging principles of regulatory evolution. Proceedings of the National Academy of Sciences of the United States of America, 104, 8605–8612.

Reed, R. D. 2004. Evidence for Notch-mediated lateral inhibition in organizing butterfly wing scales. Dev Genes Evol, 214, 43–6.

Reed, R. D., Papa, R., Martin, A., Hines, H. M., Counterman, B. A., Pardo-Diaz, C., Jiggins, C. D., Chamberlain, N. L., Kronforst, M. R., Chen, R., Halder, G., Nijhout, H. F. & Mcmillan, W. O. 2011. optix drives the repeated convergent evolution of butterfly wing pattern mimicry. Science, 333, 1137–41.

Reed, R. D. & Serf As, M. S. 2004. Butterfly wing pattern evolution is associated with changes in a Notch/Distal-less temporal pattern formation process. CurrBiol, 14, 1159–66.

Rogers, W. A., Salomone, J. R., Tacy, D. J., Camino, E. M., Davis, K. A., Rebeiz, M. & Williams, T. M. 2013. Recurrent modification of a conserved *cis*-regulatory element underlies fruit fly pigmentation diversity. PLoS Genet, 9, el003740.

Rutledge, W., Gibson, R., Siegel, E., Duke, K., Jones, R., Rucinski, D., Nunn, G., Torrence, W. A., Lewellen-Williams, C., Stewart, C., Blann, K., Belleton, L., Fincher, L., Klimberg, V. S., Greene, P., Thomas, B., Erwin, D. & Henry-Tillman, R. 2006. Arkansas Special Populations Access Network perception versus reality-cancer screening in primary care clinics. Cancer, 107, 2052–60.

Saenko, S. V., Marial Va, M. S. & Beldade, P. 2011. Involvement of the Conserved Hox gene Antennapedia in the development and evolution of a novel trait. Evodevo, 2, 9.

Sato, A., Kojima, T., Ui-Tei, K., Miyata, Y. & Saigo, K. 1999. Dfrizzled-3, a new Drosophila Wnt receptor, acting as an attenuator of Wingless signaling in wingless hypomorphic mutants. Development, 126, 4421–30.

Schilling, S., Steiner, S., Zimmerli, D. & Basler, K. 2014. A regulatory receptor network directs the range and output of the Wingless signal. Development, 141, 2483–93.

Stoehr, A. M., Walker, J. F. & Monteiro, A. 2013. Spalt expression And the development of melanic color patterns in pierid butterflies. Evodevo, 4, 6.

Treisman, J. E., Lai, Z. C. & Rubin, G. M. 1995. Shortsighted acts in the decapentaplegic pathway in Drosophila eye development and has homology to a mouse TGF-beta-responsive gene. Development, 121, 2835–45.

Urness, L. D. & Thummel, C. S. 1995. Molecular analysis of a steroid-induced regulatory hierarchy: the Drosophila E74A protein directly regulates L71-6 transcription. EMBO J, 14, 6239–46.

Van Belleghem, S. M., Rastas, P., Papanicolaou, A., Martin, S. H., Arias, C. F., Supple, M. A., Hanly, J. J., Mallet, J., Lewis, J. J., Hines, H. M., Ruiz, M., Salazar, C., Linares, M., Moreira, G. R. P., Jiggins, C. D., Counterman, B. A., Mcmillan, W. O. & Papa, R. 2017. Complex modular architecture around a simple toolkit of wing pattern genes. Nat Ecol Evol, 1.

Wallbank, R. W., Baxter, S. W., Pardo-Diaz, C., Hanly, J. J., Martin, S. H., Mallet, J., Dasmahapatra, K. K., Salazar, C., Joron, M., Nadeau, N., Mcmillan, W. O. & Jiggins, C. D. 2016. Evolutionary Novelty in a Butterfly Wing Pattern through Enhancer Shuffling. PLoS Biol, 14, el002353.

Walters, J. R., Hardcastle, T. J. & Jiggins, C. D. 2015. Sex Chromosome Dosage Compensation in Heliconius Butterflies: Global yet Still Incomplete? Genome Biol Evol, 7, 2545–59.

Warren, R. W., Nagy, L., Selegue, J., Gates, J. & Carroll, S. 1994. Evolution Of Homeotic gene regulation and function in flies and butterflies. Nature, 372, 458–61.

Weatherbee, S. D., Nijhout, H. F., Grunert, L. W., Halder, G., Galant, R., Selegue, J. & Carroll, S. 1999. Ultrabithorax function in butterfly wings and the evolution of insect wing patterns. CurrBiol, 9, 109–15.

Yassin, A., Delaney, E. K., Reddiex, A. J., Seher, T. D., Bastide, H., Appleton, N. C., Lack, J. B., David, J. R., Chenoweth, S. F., Pool, J. E. & Kopp, A. 2016. The pdm3 Locus Is a Hotspot for Recurrent Evolution of Female-Limited Color Dimorphism in Drosophila. Curr Biol, 26, 2412–2422.

Zhang, L. & Reed, R. D. 2016. Genome editing in butterflies reveals that spalt promotes and Distal-less represses eyespot colour patterns. Nat Commun, 7, 11769.

Zhang, L. L., Mazo-Vargas, A. & Reed, R. D. 2017. Single master regulatory gene coordinates the evolution and development of butterfly color and iridescence. Proceedings of the National Academy of Sciences of the United States of America, 114, 10707–10712.

