## Supplementary material for "Conservation and flexibility in the gene regulatory landscape of Heliconiine butterfly wings"

**Supplementary table 1:** number of reads per sample, % of reads which failed to map in each sample.

| *H. melpomene* | | |
| --- | --- | --- |
| Sample | Read count | % reads that fail to map |
| 13F_FP1 | 10342965 | 10.90 |
| 13G_FM1 | 14953963 | 9.56 |
| 13H_FD1 | 10205562 | 9.99 |
| 13I_HA1 | 11397631 | 13.31 |
| 13J_HPo1 | 12267336 | 11.93 |
| 14A_FP1 | 14892190 | 14.16 |
| 14B_FM1 | 12295189 | 12.44 |
| 14C_FD1 | 11528254 | 14.92 |
| 14D_HA1 | 10717086 | 15.46 |
| 14E_HPo1 | 15162459 | 12.37 |
| 14F_FP1 | 12040114 | 13.71 |
| 14G_FM1 | 12411856 | 12.78 |
| 14H_FD1 | 12303691 | 11.12 |
| 24D_HA1 | 13131538 | 32.69 |
| 14J_HPo1 | 13945427 | 14.40 |
| 15A_FP2. | 11437808 | 13.87 |
| 15B_FM2. | 10993529 | 12.81 |
| 15C_FD2. | 11856974 | 13.56 |
| 15D_HA2. | 10581808 | 13.95 |
| 15E_HPo2 | 13681765 | 11.02 |
| 16A_FP2. | 14736312 | 8.84 |
| 16B_FM2. | 13626298 | 9.40 |
| 16D_HA2. | 12686225 | 4.96 |
| 16E_HPo2 | 12823781 | 9.95 |
| 16H_FD2. | 15598458 | 10.46 |
| 17F_FP2. | 11015917 | 7.64 |
| 17G_FM2. | 12136669 | 7.46 |
| 17H_FD2. | 14589976 | 7.52 |
| 17I_HA2. | 14663720 | 7.54 |
| 17J_HPo2 | 14600239 | 7.81 |

| *H. erato* | | |
| --- | --- | --- |
| Sample | Read count | % reads that fail to map |
| B8_FP1 | 13286804 | 7.56 |
| C3_FD1 | 15104587 | 9.84 |
| C6_HA1 | 12841432 | 6.31 |
| D10_FP1 | 15919485 | 8.05 |
| F21_HPo1 | 14646293 | 8.35 |
| F8_FM1 | 14965698 | 10.61 |
| G21_FM1 | 13046272 | 8.03 |
| G5_FM1 | 12543875 | 7.66 |
| H4_HA1 | 13419952 | 7.91 |
| I8_FD1 | 14022680 | 10.02 |
| S3_HPo1 | 24574848 | 6.98 |
| A4_HPo2 | 12912895 | 18.97 |
| B2_FD2 | 13435820 | 19.43 |
| B4_HA2 | 12182847 | 18.86 |
| C4_FM2 | 14127156 | 20.68 |
| C9_FP2 | 15100384 | 19.49 |
| D6_HA2 | 14027838 | 17.72 |
| D9_HPo2 | 12354324 | 17.87 |
| E3_FM2 | 13926825 | 23.33 |
| E6_FP2 | 13622452 | 15.73 |
| G6_FD2 | 13281517 | 19.52 |
| J4_FD1 | 14498546 | 16.59 |
| J8_FP2 | 16882308 | 19.12 |
| N3_FM2 | 13139640 | 18.78 |

| *Agraulis vanillae* | | |
| --- | --- | --- |
| Sample | Read count | % reads that fail to map |
| 11F | 21601472 | 10.78 |
| 11G | 19505732 | 10.39 |
| 11H | 9472553 | 11.65 |
| 11I | 20695225 | 16.69 |
| 11J | 15779259 | 12.80 |
| 12A | 17909757 | 18.40 |
| 12B | 21362299 | 9.44 |
| 12C | 19020085 | 10.37 |
| 12D | 21698478 | 11.32 |
| 12E | 21574847 | 11.73 |
| 12F | 19511174 | 10.86 |
| 12G | 23853328 | 11.20 |
| 12H | 23137900 | 10.96 |
| 12I | 18912480 | 12.82 |
| 12J | 18012267 | 13.16 |
| 21A | 25105152 | 11.28 |
| 21D | 21967288 | 11.08 |
| 21E | 22224755 | 12.23 |
| 22G | 18790997 | 7.53 |
| 22H | 19159884 | 7.12 |
| 23A | 16849671 | 8.19 |
| 23B | 22852458 | 7.84 |
| 23C | 20814817 | 7.90 |
| 23D | 21198748 | 7.27 |
| 23E | 21347562 | 7.28 |
| 23F | 22977885 | 8.09 |
| 23G | 19254249 | 7.79 |
| 23H | 19779195 | 7.88 |
| 23I | 20971110 | 8.80 |
| 23J | 23057419 | 7.51 |

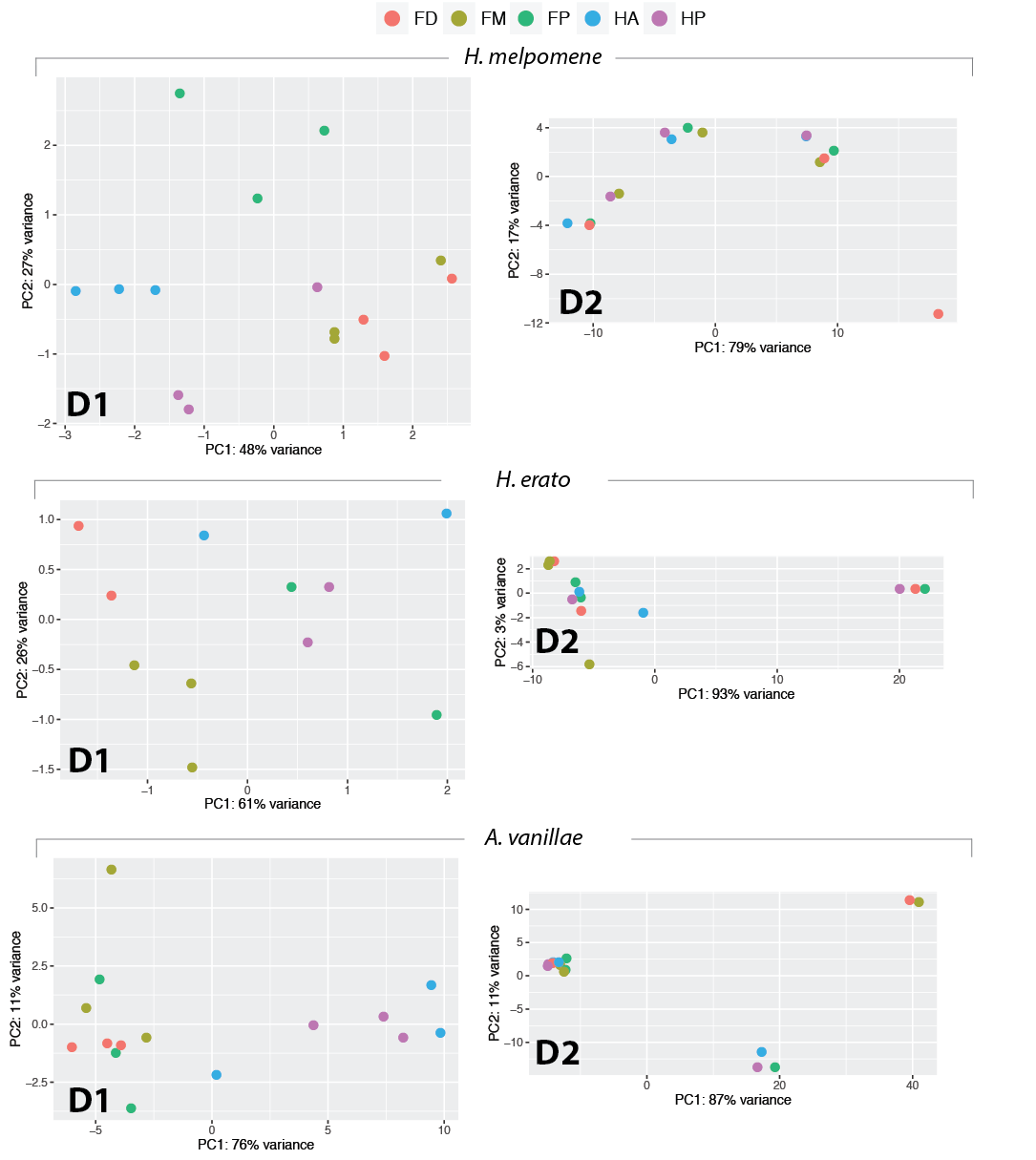

**Supplementary figure 1:** Principle component analysis of RNAseq data. Each point corresponds to a sample, with colours corresponding to wing compartment. (FP – proximal forewing, FM – medial forewing, FD – distal forewing, HA – anterior forewing, HP – posterior forewing.

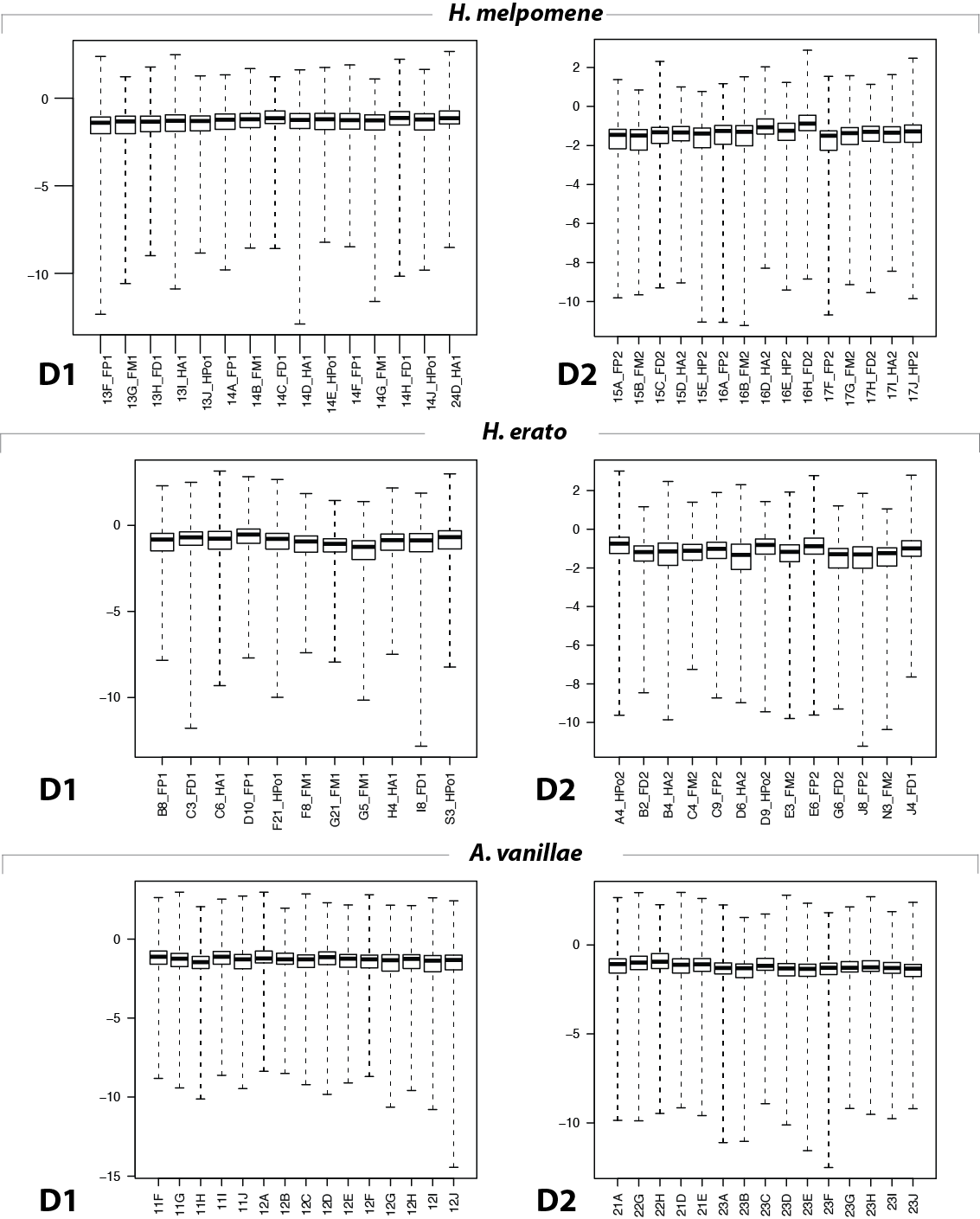

**Supplementary figure 2:** Cook’s distance for each sample.

Supplementary table 2: genes differentially expressed in larval wing discs of *H. erato* and *H. erato.*

| Genes upregulated in forewings | | |
| --- | --- | --- |
| *nov_gene_002001* |  |  |
| *ham* | hamlet | neuron fate selection, type II neuroblast maturation |
| *AOX3* | Aldehyde oxidase 3 |  |
| *Dhpr* | Dihydropteridine reductase |  |
| *nov_gene007136* |  |  |
| *PK2-R1* | Pyrokinin 2 receptor 1 | GPCR, binds Hug (an NT and chitin synthesis regulator) |
| *nov_gene009808* |  |  |
| *CG10298* | no annotated func |  |
| *nov_gene034094* |  |  |
| *nov_gene034095* |  |  |
| *Pepck* | Phosphoenolpyruvate carboxykinase |  |
| *nov_gene035885* |  |  |
| *nov_gene036476* |  |  |
| *Nrk* | Neurospecific receptor kinase | axon pathfinding, rhabdomeere elongation |
| *nov_gene045731* |  |  |
| Genes upregulated in hindwings | | |
| *nrv2* | nervana 2 | Na/K ATPase subunit 2. |
| *Cht2* | Chitinase 2 |  |
| *AdamTS-A* | ADAM metallopeptidase with thrombospondin type 1 motif A | secreted matrix metalloprotease (mutants show apical surface irregularities). |
| *CG3168* |  | anion transmembrane transporter |
| *Ubx* | Ultrabithorax | homeodomain transcription factor |
| *pigs* | pickled eggs | microtubule binding, negative regulator of notch |
| Genes differentially regulated in both species, but in the opposite direction | | |
| *r-l* | rudimentary-like | orotate phosphoribosyltransferase |
| *nov_gene013098* |  |  |

**Supplementary table 3:**

| Gene |  |  |
| --- | --- | --- |
| nov_gene015670 |  |  |
| Hr38 | Hormone receptor-like in 38 | required for adult cuticle development |
| gcm | glial cells missing | (discussed in text) |
| Sse | Separase | chromatid separation in meiosis |
| sand | sandman | dopamine responsive potassium channel |
| CG2663 |  | intracellular transport |
| gd | gastrulation-defective | secreted serine protease, activates Toll ligand spatzle, dorsoventral axis patterning |
| nov_gene011683 |  |  |
| CG42339 |  | immune response |
| GstS1 | Glutathione S transferase S1 |  |
| CGstD1 | Glutathione S transferase D1 |  |
| CG4914 |  | serine endopeptidase activity |
| CG7896 | no annotated func |  |
| Scp2 | Sarcoplasmic calcium-binding protein |  |
| disco-r | disco-related | Epithelial-mesenchymal transition, wing expansion |
| sob | sister of odd and bowl | leg disc – joint morphogenesis |
| Reck | Reversion-inducing-cysteine-rich protein with kazal motifs | no annotated func |
| nov_gene021747 |  |  |
| Faa | Fumarylacetoacetase | aromatic amino acid metabolism |
| Thor | Thor | translation initiation factor 4E binding protein |
| CG14257 | no annotated func |  |
| spo | spook | CNS development, cuticle development, head involuiton |
| CG42346 | no annotated func |  |
| nov_gene007135 |  |  |
| nov_gene007136 |  |  |
| nov_gene046315 |  |  |
| Cyp6a2 | Cytochrome P450-6a2 | response to DDT, caffeine. |
| zfh2 | Zn finger homeodomain 2 | putative TF, proximal-distal patterning during wing and leg imaginal disc development |
| CG4945 |  | mesoderm development |
| knrl | knirps-like | orphan nuclear hormone receptor, target of Hh, Wnt and Notch pathways |
| hth | homothoroax |  |
| nov_gene015709 |  |  |
| Cht2 | Chitinase 2 |  |
| nov_gene008541 |  |  |
| Roc1a | Regulator of cullins 1a | E3 Ubq ligase component |
| CG9380 |  |  |
| CG41378 |  |  |
| Rap2l | Ras-associated protein 2-like |  |
| scramb1 | scramblase 1 | scramble phospholipids across bilayer. (role in neurotransmission, apoptosis) |
| nov_gene031883 |  |  |
| nov_gene031884 |  |  |
| pnut | peanut | component of septin GTPase complex, involved in organization of cell cortex |
| alpha-Catr | α-catenin related | actin binding |
| nov_gene |  |  |
| CG9701 | beta-glucosidase |  |
| CG11370 | no annotated func |  |
| CG15239 | no annotated func |  |
| sqd | squid | RNA binding protein, localisation |
| Nlg3 | Neuroligin 3 | synaptic adhesion molecule involved in synapse formation and synaptic transmission |
| Cyp6a18 |  |  |
| CG15497 | no annotated func |  |
| E2f1 | E2F transcription factor 1 |  |
| Archease |  | axon regenration |
| DCTN3-p24 | Dynactin 3, p24 subunit | dynein activation |
| Fip1 | Factor interacting with poly(A) polymerase 1 |  |
| pum | pumilio | 3’UTR binding protein |
| pum |  |  |
| mago | mago nashi | splicing, photoreceptor differentiation |
| RpS8 | Ribosomal protein S8 |  |
| Spn85F | Serpin 85F | chitin modifier |
| Mctp | Multiple C2 domain and transmembrane region protein | Ca^2+^ ion-binding, membrane component |
| KFase | Kynurenine formamidase | possible pigment synthesis! |
| obst-A | obstructor-A | chitin maturation |
| obst-A |  |  |
| obst-B |  |  |
| betaTub56D | β-Tubulin at 56D | microtubule unit |
| CG3655 | no annotated func |  |
| CG4678 |  | metallocarboxypeptidase activity |
| CG34461 |  | chitin strucutural component |
| nov_gene034193 |  |  |
| Lcp65Ac | Lcp65Ac | chitin strucutural component |
| Cpr49Aa | Cuticular protein 49Aa | chitin strucutural component |
| nov_gene034334 |  |  |
| unc-5 | unc-5 | axon guidance |
| Nc73EF | Neural conserved at 73EF |  |
| nov_gene036907 |  |  |
| Spn42Da | Serpin 42Da |  |
| nov_gene046015 |  |  |
| beat-IV | beat-IV | heterophilic cell-cell adhesion |
| nGstE6 |  |  |
| CG42259 |  | haemolymph coagulation |
| CG42259 |  |  |
| nov_gene007986 |  |  |
| CG1402 | no annotated func |  |
| Treh | Trehalase | trehalose metabooism, neural stem cell maintenance |
| Chd64 |  | juvenile hormone-responsive, actin binding |
| CG4213 | no annotated func |  |
| H2.0 | Homeodomain protein 2.0 |  |
| CG16885 | no annotated func |  |
| IP3K2 | IP3-kinase 2 | regulates calcium levels by influencing IP3 signaling |
| nov_gene011713 |  |  |
| CG16786 | no annotated func |  |
| Bx | Beadex | dimerizes with LIM transcription factors including Apterous in the wing disc. |
| Ets98B |  | oocyte-expressed |
| neur | neuralized | endocytosis-dependent activation of Notch |
| CG42390 | no annotated func |  |
| Nrg | Neuroglian | axon growth, imaginal disc morphognesis |
| CG40160 |  | serine-type endopeptidase |
| CG8369 |  | wing disc dorsal-ventral patterning (via interaction with Ap-Bx) |
| CG10407 | no annotated func |  |
| Toll-7 |  | axon guidance, viral immunity |
| mspo | M-spondin | regulation of myoblast fusion |
| nov_gene039499 |  |  |
| ds | dachsous | cadherin binding, Planar Cell Polarity |
| beat-IIIb |  | heterophilic cell-cell adhesion |
| Nos | Nitric oxide synthase | nervous system development |
| TwdlE | TweedleE | chitin strucutural component |
| CG14964 |  | actin binding protein |
| Prm | Paramyosin | structural component of muscle |

Supplementary table 4

| gene | day 2 |  |
| --- | --- | --- |
| DE in *Agraulis* & *H. erato* | | |
| CG13367 |  |  |
| nov_gene007134 |  |  |
| Dnai2 | dynein, axonemal, intermediate chain 2 |  |
| nov_gene00837 |  |  |
| nov_gene008541 |  |  |
| ple | pale | tyrosine hydroxylase (rate limiting step in melanin synthesis) |
| Cpr65Ec | Cuticular protein 65Ec | chitin structural component |
| CG10555 |  | transcription coactivator |
| Ddc | Dopa decarboxylase | melanin synthesis |
| CG33290 | no annotated func |  |
| Prm | Paramyosin | structural component of muscle |
| Mlc1 | Myosin alkali light chain 1 | myosin component |
| CG9297 | no annotated func |  |
| CG11825 | no annotated func |  |
| DE in *H. melpomene* & *H. erato* | | |
| nov_gene011805 |  |  |
| mab-21 | malformed abdomen 21 | no annotated func |
| Sulf1 | Sulfated | heparan sulfate modifier, regulates Wnt ligand diffusion, als hh signalling |
| gd | gastrulation-defective | secreted serine protease, activates Toll ligand spatzle, dorsoventral axis patterning |
| t | tan | converts NBAD to dopamine – melanin synthesis pathway |
| CG4757 |  | carboxylic ester hydroxylase |
| CG5973 |  | transporter activity |
| AP-2alpha | Adaptor Protein complex 2, α subunit | cell transport |
| DE in *H. melpomene* & *Agraulis* | | |
| CG33290 | no annotated func |  |
| CG31954 |  | serine endopeptidase |
| nov_gene011684 |  |  |
| CG5112 | no annotated func |  |
| Mctp | Multiple C2 domain and transmembrane region protein |  |
| nov_gene037141 |  |  |
| CG13868 | no annotated func |  |
| nov_gene046277 |  |  |
| nov_gene007986 |  |  |

Supplementary table 5: Wnt pathway constituents and gene codes

| gene_code | gene_type | *H. melpomene* | *H. erato* |  |
| --- | --- | --- | --- | --- |
| fz | receptor | HMEL030072 | evm.model.Herato0101.604 | Gene.17645::c50081_g1_i1::g.17645::m.17645 evm.model.Herato0101.604 |
| Wg | ligand | HMEL022601 | evm.model.Herato0101.738 | Gene.28080::c53180_g1_i1::g.28080::m.28080 evm.model.Herato0101.735 |
| Wnt10 | ligand | HMEL011434 | evm.model.Herato0101.735 | Gene.22452::c51642_g1_i1::g.22452::m.22452 evm.model.Herato0101.736 |
| Wnt6 | ligand | HMEL011436 | evm.model.Herato0101.736 | Gene.24659::c52260_g1_i1::g.24659::m.24659 evm.model.Herato0101.738 |
| Wnt4 | ligand | HMEL011441 | evm.model.Herato0101.740 | Gene.44206::c56654_g1_i1::g.44206::m.44206 evm.model.Herato0101.740 |
| VhaM8.9 | PCP | HMEL043115 | evm.model.Herato0310.108 | Gene.14196::c48712_g1_i1::g.14196::m.14196 evm.model.Herato0310.108 |
| Groucho | TF | HMEL016524 | evm.model.Herato0403.3 | Gene.19315::c50670_g3_i1::g.19315::m.19315 evm.model.Herato0403.3 |
| pygo | TF | HMEL004050 | evm.model.Herato0405.22 | Gene.36671::c55176_g1_i1::g.36671::m.36671 evm.model.Herato0405.22 |
| gilgamesh | intracellular | HMEL043699 | evm.model.Herato0405.46 | Gene.9988::c46287_g1_i1::g.9988::m.9988 evm.model.Herato0405.46 |
| spen_1 | TF | HMEL042399 | evm.model.Herato0503.212 |  |
| Rho1 | intracellular | HMEL016025 | evm.model.Herato0503.23 | Gene.22721::c51732_g1_i1::g.22721::m.22721 evm.model.Herato0503.212 |
| fz3 | receptor | HMEL044184 | evm.model.Herato0503.9 | Gene.35454::c54867_g1_i1::g.35454::m.35454 evm.model.Herato0503.23 |
| CG8786 | intracellular | HMEL046268 | evm.model.Herato0701.527 | Gene.31107::c53893_g1_i1::g.31107::m.31107 evm.model.Herato0503.9 |
| fz2_1 | receptor | HMEL046532 | evm.model.Herato0701.759 | Gene.17796::c50150_g1_i1::g.17796::m.17796 evm.model.Herato0701.527 |
| osa | TF | HMEL030920 | evm.model.Herato0901.275 | Gene.26847::c52848_g1_i1::g.26847::m.26847 |
| Wnt5 | ligand | HMEL018100 | evm.model.Herato1001.160 | Gene.22486::c51656_g2_i1::g.22486::m.22486 evm.model.Herato0901.275 |
| WntA | ligand | HMEL018100 | evm.model.Herato1001.161 | Gene.29008::c53376_g1_i6::g.29008::m.29008 evm.model.Herato1001.160 |
| Wnt2 | ligand | HMEL022591 | evm.model.Herato1001.69 | Gene.29005::c53376_g1_i1::g.29005::m.29005 evm.model.Herato1001.161 |
| Wnt5b | ligand | HMEL022606 | evm.model.Herato1001.88 | Gene.14793::c48962_g1_i1::g.14793::m.14793 evm.model.Herato1001.69 |
| shf | heparan_interacting | HMEL031574 | evm.model.Herato1003.133 | Gene.24659::c52260_g1_i1::g.24659::m.24659 evm.model.Herato1001.88 |
| ttv | heparan_interacting | HMEL013339 | evm.model.Herato1005.143 | Gene.9275::c45698_g2_i1::g.9275::m.9275 evm.model.Herato1003.133 |
| Drl-2 | ligand | HMEL015485 | evm.model.Herato1007.90 | Gene.19220::c50654_g1_i1::g.19220::m.19220 evm.model.Herato1005.143 |
| dally | intracellular | HMEL015252 | evm.model.Herato1108.113 | Gene.48364::c57401_g1_i1::g.48364::m.48364 evm.model.Herato1007.90 |
| botv | heparan_interacting | HMEL011746 | evm.model.Herato1108.259 | Gene.37419::c55339_g2_i1::g.37419::m.37419 evm.model.Herato1108.113 |
| dlp | intracellular | HMEL032267 | evm.model.Herato1108.30 | Gene.46191::c57019_g1_i1::g.46191::m.46191 evm.model.Herato1108.259 |
| hyrax | TF | HMEL015680 | evm.model.Herato1108.345 | Gene.38015::c55477_g2_i1::g.38015::m.38015 evm.model.Herato1108.30 |
| boca | intracellular | HMEL033550 | evm.model.Herato1202.553 | Gene.44686::c56735_g1_i1::g.44686::m.44686 evm.model.Herato1108.345 |
| dx | intracellular | HMEL007293 | evm.model.Herato1202.820 | Gene.38286::c55532_g1_i1::g.38286 |
| sfl | heparan_interacting | HMEL034074 | evm.model.Herato1301.128 | Gene.19786::c50823_g1_i2::g.19786::m.19786 evm.model.Herato1202.820 |
| kermit | intracellular | HMEL034176 | evm.model.Herato1301.227 | Gene.21881::c51480_g1_i1::g.21881::m.21881 evm.model.Herato1301.128 |
| stan | PCP | HMEL034041 | evm.model.Herato1301.96 | Gene.16962::c49871_g1_i1::g.16962::m.16962 evm.model.Herato1301.227 |
| Pontin | TF | HMEL017143 | evm.model.Herato1301.391 | Gene.3686::c33770_g1_i1::g.3686::m.3686 evm.model.Herato1301.96 |
| alphao | intracellular | HMEL035171 | evm.model.Herato1408.158 | Gene.19767::c50819_g1_i1::g.19767::m.19767 evm.model.Herato1301.391 |
| mwh | PCP | HMEL035679 | evm.model.Herato1505.49 | Gene.22108::c51540_g4_i2::g.22108::m.22108 evm.model.Herato1408.158 |
| Axin | PCP | HMEL036070 | evm.model.Herato1507.286 | Gene.32473::c54230_g1_i2::g.32473::m.32473 evm.model.Herato1505.49 |
| dco | intracellular | HMEL035880 | evm.model.Herato1507.83 | Gene.46731::c57131_g1_i2::g.46731::m.46731 evm.model.Herato1507.286 |
| sgg | intracellular | HMEL012308 | evm.model.Herato1603.51 | Gene.39804::c55818_g2_i1::g.39804::m.39804 evm.model.Herato1507.83 |
| nejire | TF | HMEL038398 | evm.model.Herato1703.13 | Gene.4876::c38729_g2_i1::g.4876::m.4876 evm.model.Herato1603.51 |
| dishevelled | intracellular | HMEL038512 | evm.model.Herato1705.40 | Gene.13733::c48458_g1_i1::g.13733::m.13733 evm.model.Herato1703.13 |
| sgl | intracellular | HMEL010174 | evm.model.Herato1705.41 | Gene.21729::c51423_g2_i1::g.21729::m.21729 evm.model.Herato1705.40 |
| Ext2 | heparan_interacting | HMEL017833 | evm.model.Herato1805.156 | Gene.38119::c55499_g1_i1::g.38119::m.38119 evm.model.Herato1705.41 |
| Notum | heparan_interacting | HMEL039208 | evm.model.Herato1805.28 | Gene.43087::c56437_g1_i1::g.43087::m.43087 evm.model.Herato1805.156 |
| Rfabg | heparan_interacting | HMEL002964 | evm.model.Herato1807.120 | Gene.12118::c47616_g1_i1::g.12118::m.12118 evm.model.Herato1805.28 |
| pangolin | TF | HMEL039657 | evm.model.Herato1807.48 | Gene.1076::c12781_g1_i1::g.1076::m.1076 evm.model.Herato1807.120 |
| CkIalpha | intracellular | HMEL016957 | evm.model.Herato1901.49 | Gene.30555::c53776_g1_i2::g.30555::m.30555 evm.model.Herato1807.48 |
| CtBP | intracellular | HMEL016973 | evm.model.Herato1901.62 | Gene.37459::c55348_g3_i1::g.37459::m.37459 evm.model.Herato1901.49 |
| twins | intracellular | HMEL013172 | evm.model.Herato1904.66 | Gene.11911::c47529_g1_i1::g.11911::m.11911 evm.model.Herato1901.62 |
| arm_B-Catenin | intracellular | HMEL008024 | evm.model.Herato2001.403 | Gene.15402::c49262_g1_i1::g.15402::m.15402 evm.model.Herato1904.66 |
| fry | intracellular | HMEL011016 | evm.model.Herato2101.61_evm.model.Herato2101.59 | Gene.40676::c56009_g1_i1::g.40676::m.40676 evm.model.Herato2001.403 |
| wntless | intracellular | HMEL032988 | novel_model_155 | Gene.47366::c57232_g1_i2::g.47366::m.47366 evm.model.Herato2101.61_evm.model.Herato2101.59 |
|  |  |  |  | Gene.15032::c49092_g1_i1::g.15032::m.15032 novel_model_155 |

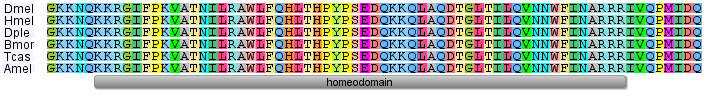

**Supplementary Figure S3. Homothorax homeodomain protein sequence is completely conserved between insects.** Top to bottom: *Drosophila melanogaster, Heliconius melpomene, Danaus plexippus, Bombyx mori, Tribolium castaneum, Apis mellifera.* Only homeodomain is shown, there is lower conservation outside this domain.

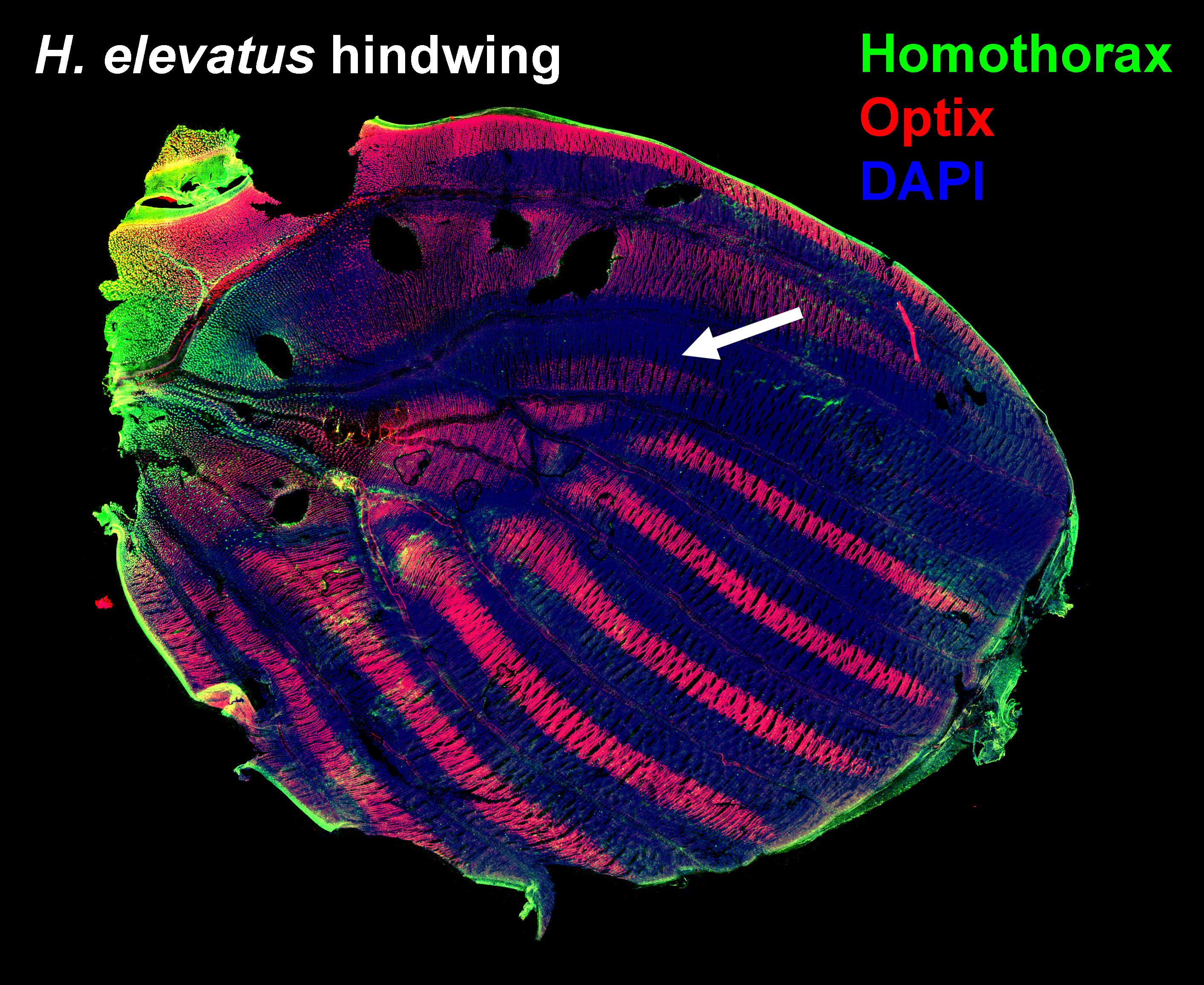

**Supplementary Figure S5. Homothorax and Optix expression in the hindwing.**

In both *H. elevatus* and *H. rosina* hindwings Homothorax forms a limited gradient in the hinge region. This does not correspond to the *dennis* hindwing bar (arrow), suggesting this part of the pattern is influenced by factors other than Homothorax.

60-72 hrs post-pupation *H. elevatus*.
